# Piezo1 Triggers an Angiopoietin-2-Integrin Signaling Loop in Schlemm’s Canal to Regulate Intraocular Pressure

**DOI:** 10.1101/2025.10.24.683742

**Authors:** Naoki Kiyota, Dilip K. Deb, Hongyuan Ren, Kerolous Salama, Yalu Zhou, Haiyan Gong, Benjamin R. Thomson, Susan E. Quaggin

## Abstract

Elevated intraocular pressure (IOP), driven by increased outflow resistance in the trabecular meshwork and Schlemm’s canal (SC), is a primary risk factor for glaucoma. This resistance is regulated by broadly active endothelial signaling systems such as ANGPT1-TIE2 and by dynamic flow-responsive pathways that remain poorly understood. Here, we identify a previously unrecognized, TIE2-independent, mechanosensitive ANGPT2–integrin α9β1 pathway in the SC endothelium that regulates IOP. In vitro and in vivo, we show that activation of the mechanosensory channel PIEZO1 triggers ANGPT2 secretion and promotes cell-surface clustering of integrin α9β1. Deletion of SC-expressed *Piezo1* or *Itga9* in mice resulted in SC narrowing, impaired flow-mediated SC endothelial proliferation, IOP elevation and glaucoma. Furthermore, ANGPT2 deficiency or blockade disrupted PIEZO1-induced integrin activation and reduced aqueous humor outflow facility. These findings establish autocrine PIEZO1–ANGPT2–ITGA9 signaling as a link between mechanosensory stimuli, SC structure and IOP regulation, offering promising new targets for glaucoma therapy.

---

Glaucoma is a leading cause of blindness, affecting ∼64 million people worldwide^1^. While several distinct forms of glaucoma exist^2^, loss of retinal ganglion cells (RGCs) and deformation of the optic nerve are directly responsible for vision loss in all forms of the disease. While glaucomatous damage occurs in the posterior segment, elevated intraocular pressure (IOP) due to increased aqueous humor outflow (AHO) resistance^3,4^ is the primary risk factor for pathogenesis and IOP reduction slows vision loss in both high-pressure and normotensive glaucoma^5,6^. Schlemm’s canal (SC), a large vessel in the ocular anterior segment, is responsible for the majority of AHO, and alterations in physical characteristics of SC endothelial cells are associated with increased outflow resistance and elevated IOP, making them a key therapeutic target^7,8^.

SC is a unique endothelium, expressing a hybrid phenotype with both venous and lymphatic characteristics^9,10^. Consistent with its role in modulating IOP and AHO, genes regulating SC development and function have been linked with glaucoma in patients and animal models. Some of the best characterized belong to the angiopoietin (ANGPT) signaling pathway, comprised of the ANGPT ligands and the SC-expressed receptor TIE2 (encoded by *TEK*)^11,12^.

Loss-of-function variants in TEK or its trabecular meshwork (TM)-expressed ligand ANGPT1 cause primary congenital glaucoma (PCG) in humans and mice and are associated with increased risk of primary open-angle glaucoma (POAG), the most common form of the disease^11–13^. In addition to this ANGPT1-mediated paracrine signaling pathway, a second ANGPT ligand, ANGPT2, is expressed by SC endothelial cells themselves and acts in an autocrine fashion^12^. Mouse studies have suggested ANGPT2 is less important than paracrine ANGPT1 for canal development, but *ANGPT2* variants have been associated with POAG^14^ and a noncoding *ANGPT2* variant that increased *ANGPT2* expression and SC size in mice has been associated with reduced POAG risk, suggesting ANGPT2 has a significant functional role^15^. Furthermore, compound deletion of *Angpt1* and *Angpt2* caused more severe SC attenuation in mice than *Angpt1* deletion alone, indicating compensation by ANGPT2^12^.

In addition to its role in TIE2 signaling, ANGPT2 has TIE2-independent roles in other vascular beds, functioning through integrins to regulate endothelial permeability, migration, and cell proliferation^16^. In lymphatic endothelial cells (LECs), which share key features with SC, autocrine ANGPT2–TIE2 signaling activated by the mechanosensitive Ca^2+^ channel PIEZO1 enables LECs to transduce shear stress into TIE2-dependent molecular and functional responses^17,18^. Like lymphatic vessels, SC is regulated by mechanical stimuli and shear stress^19^, and as *Piezo1* regulates IOP in mice and *PIEZO1* variants have been linked to POAG, we speculated that PIEZO1 plays a similar role in SC^20,21^. Here, we interrogate this pathway in detail and show that within SC endothelial cells, PIEZO1 activation induces ANGPT2 secretion, leading to TIE2 phosphorylation and downstream signaling. In addition, this ANGPT2 secretion regulates integrin signaling in a TIE2-independent manner through endothelial integrin α9β1 and phosphorylation of focal adhesion kinase (FAK). Together, these findings provide new evidence for the importance of non-canonical angiopoietin signaling within the canal endothelium and identify new targets for future therapeutic development.

## Results

We have previously reported that PIEZO1 activation in primary cultured LECs initiates a mechanotransduction cascade involving ANGPT2 secretion, TIE2 phosphorylation, and nuclear export of the transcription factor FOXO1, ultimately upregulating lymphatic valve–associated genes including *GJA4* and *ITGA9*^17^. Components of this pathway are detected in tissues of the iridocorneal angle, and human genetic studies have implicated variants—including rare coding variants—in *PIEZO1* and *TEK*, and loci at/near *ANGPT2* and *SVEP1* (an extracellular matrix ligand for ITGA9) in IOP regulation (and thus ocular hypertension) and/or primary open-angle glaucoma (POAG) risk^15,21–25^. However, while SC endothelial cells exhibit aspects of lymphatic endothelial fate, they are distinct from true lymphatics and retain features of their original blood-vascular phenotype^9,10^, along with adaptations to their unique physiological environment. To determine whether a similar PIEZO1-dependent ANGPT2–TIE2 signaling axis exists in SC and regulates AHO, we first confirmed that all members of the signaling cascade were expressed in SC endothelial cells. Prior studies have reported high levels of ANGPT2, ITGA9, and TEK in SC^12,26^. Analysis of a previously published single-cell RNA-seq dataset^12,26^ revealed that although *Piezo1* expression was detected in the TM as previously reported^20,27^, expression was higher in SC endothelium (**Fig. 1A**).

**Figure 1.**
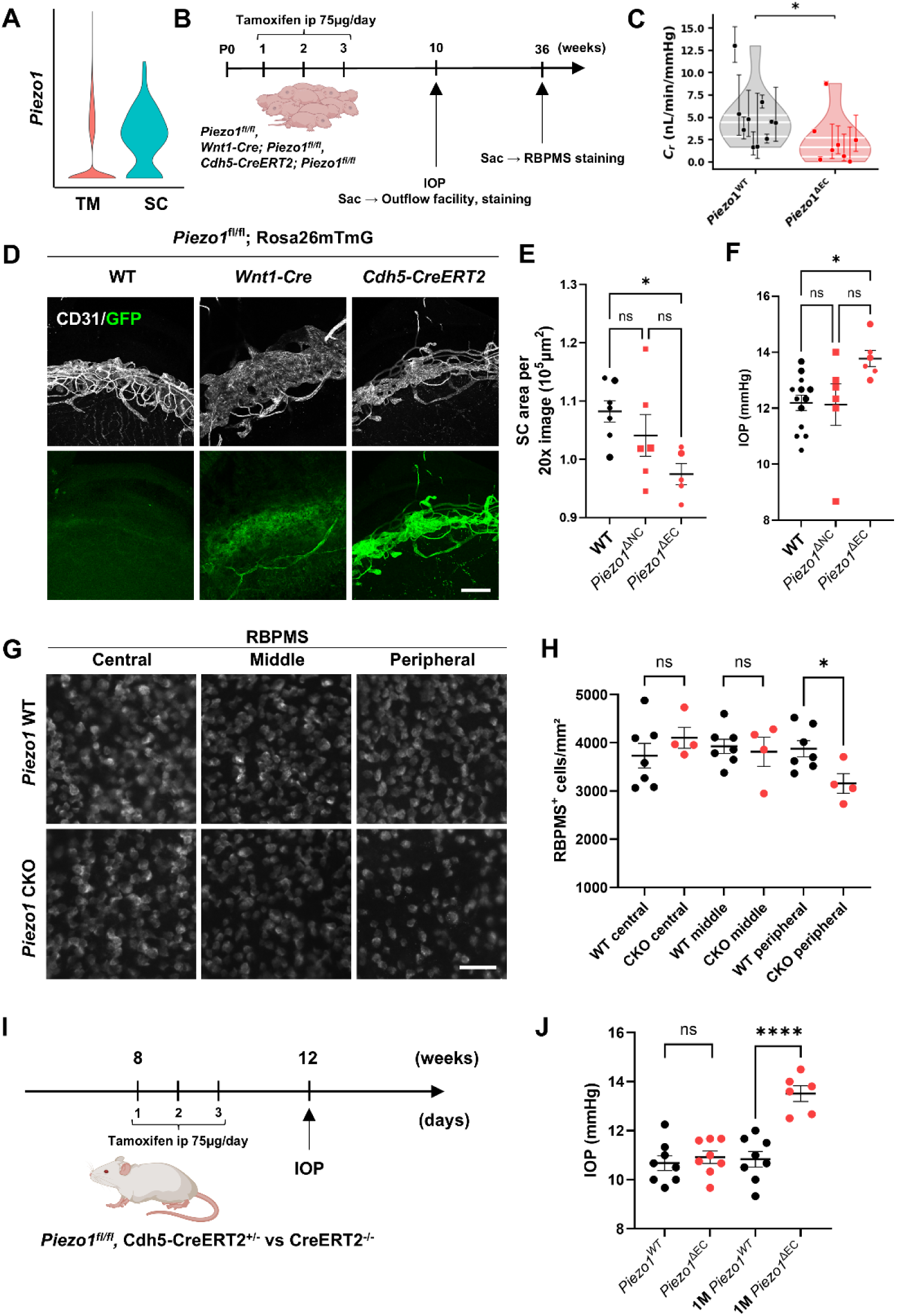
Endothelial *Piezo1* deletion reduces outflow facility, narrows SC, elevates IOP, and induces peripheral RGC loss. (A) Violin plot from single-cell RNA-seq re-analysis showing *Piezo1* expression in TM and SC endothelium. (B) Experimental timeline for neural-crest–specific (Wnt1-Cre; *Piezo1*^ΔNC^) and endothelial-specific (Cdh5-CreERT2; *Piezo1*^ΔEC^) deletion; tamoxifen (75 µg/day, i.p.) was administered at P1–P3 for the endothelial model, followed by assessment of outflow facility, SC staining, and IOP at 10 weeks and RBPMS staining at 36 weeks. (C) Outflow facility (C_r_) is reduced in *Piezo1*^ΔEC^ compared with *Piezo1* WT. (D) Recombination-site validation in limbal whole mounts crossed to Rosa26mTmG reporters showing CD31 (white) and mTmG-GFP (green), confirming TM labeling in the Wnt1-Cre line and SC-endothelial labeling in the Cdh5-CreERT2 line. (E) Quantification of SC area per 20× image shows a reduction in *Piezo1*^ΔEC^, whereas *Piezo1*^ΔNC^ does not differ from WT. (F) IOP at 10 weeks is increased in *Piezo1*^ΔEC^ relative to WT and *Piezo1*^ΔNC^. (G) Representative RBPMS-stained retinal flat mounts at 36 weeks (central, middle, and peripheral regions) from WT and *Piezo1*^ΔEC^. (H) Quantification shows a selective reduction of RGC density in the peripheral retina of *Piezo1*^ΔEC^, with central and middle regions unchanged. (I) Experimental timeline for inducible adult endothelial *Piezo1* deletion (*Piezo1*^fl/fl^; Cdh5-CreERT2^+/−^; *Piezo1*^ΔEC^). Tamoxifen (75 µg/day, i.p.) was administered once daily for three consecutive days at 8 weeks of age, and IOP was assessed at baseline and one month after induction. (J) Baseline IOP does not differ between WT and *Piezo1*^ΔEC^ mice, whereas IOP is significantly elevated in *Piezo1*^ΔEC^ relative to time-matched WT one month after induction. Statistics: two-group comparisons, unpaired two-tailed Student’s t-test; multi-group comparisons, one-way ANOVA with Tukey–Kramer post hoc test. Scale bars: (D) 100 µm; (G) 50 µm. Data are presented as mean ± SEM. *P < 0.05, **P < 0.01, ns = not significant.

### Endothelial-specific Piezo1 deletion narrows SC, reduces outflow facility, elevates IOP, and reduces peripheral RGC density

As described above, prior studies detected *Piezo1* transcripts in the TM and, by single-cell profiling, in endothelial populations of the conventional outflow pathway, including SC^20,27^. Our single-cell RNA-seq analysis^26^ likewise showed *Piezo1* expression in both TM and SC, with higher levels in SC endothelium (**Fig. 1A**). To determine the importance of *Piezo1* expression in each tissue for IOP homeostasis, we generated two parallel knockout mouse lines by crossing *Piezo1*^fl/fl^ conditional knockout mice with animals expressing either Wnt1-Cre (targeting neural crest–derived TM cells) or Cdh5-CreERT2 (targeting vascular endothelium, including SC). Hereafter, we refer to Wnt1-Cre–mediated deletion as neural-crest–specific deletion (*Piezo1*^ΔNC^) and to Cdh5-CreERT2–mediated deletion as endothelial-specific deletion (*Piezo1*^ΔEC^). Tamoxifen (75 μg/day, i.p., P1–P3) was administered to induce recombination in Cdh5-CreERT2 mice (**Fig. 1B**). Tissue specificity of each Cre-expressing strain was validated by crossing each line to Rosa26mTmG reporters for recombination site validation, confirming efficient targeting of TM by Wnt1-Cre and of SC endothelium by Cdh5-CreERT2 (**Fig. 1D**). At 10 weeks of age, we measured IOP and outflow facility, followed by SC morphometry by CD31 immunostaining. Outflow facility (Cr; reference outflow facility evaluated at Pr = 8 mmHg, see Methods) was reduced in *Piezo1*^ΔEC^ compared with *Piezo1* wild-type (WT) (2.36 ± 2.84 vs 4.84 ± 3.29 nL/min/mmHg; n = 8–10 eyes per group; P = 0.030; **Fig. 1C**). *Piezo1*^ΔEC^ mice showed a significant reduction in SC area relative to controls, whereas *Piezo1*^ΔNC^ did not differ (**Fig. 1E**; one-way ANOVA with Tukey–Kramer, n = 5–7 eyes per group; adjusted P values as indicated), indicating that endothelial PIEZO1 is required to maintain SC area. Consistent with this phenotype, IOP at 10 weeks was higher in *Piezo1*^ΔEC^ than WT (13.77 ± 0.28 vs 12.78 ± 0.25 mmHg; Tukey-adjusted P = 0.026), while *Piezo1*^ΔNC^ did not differ from WT (12.12 ± 0.75 mmHg; Tukey-adjusted P = 0.438) (**Fig. 1F**; n = 6–13 mice per group). To assess the consequences of chronic IOP elevation, we quantified RNA-binding protein with multiple splicing (RBPMS)-positive RGCs at 36 weeks in *Piezo1*^fl/fl^; Cdh5-CreERT2^−^ (control) and *Piezo1*^fl/fl^; Cdh5-CreERT2^+^ (*Piezo1*^ΔEC^) mice. Counts in the central, middle, and peripheral retina revealed a selective reduction in the peripheral region of *Piezo1*^ΔEC^ mice (**Fig. 1G,H**; unpaired two-tailed t-tests within each eccentricity; n = 4–7 eyes per group; peripheral: P < 0.05), consistent with regionally biased RGC degeneration secondary to ocular hypertension. To test whether *Piezo1* was also required for adult IOP homeostasis, we induced endothelial deletion in adult *Piezo1*^fl/fl^; Cdh5-CreERT2^+^ mice (*Piezo1*^ΔEC^) by administering tamoxifen for three consecutive days at 8 weeks of age, with Cre^−^ littermates as controls (**Fig. 1I**). Baseline IOP did not differ between genotypes (WT, 10.68 ± 0.30 vs *Piezo1*^fl/fl^; Cdh5-CreERT2^+^, 10.92 ± 0.25 mmHg; P = 0.547; n = 8 per group), whereas IOP was markedly elevated one month after induction (1M WT, 10.83 ± 0.32 vs 1M *Piezo1*^ΔEC^, 13.51 ± 0.32 mmHg; P < 0.0001; n = 6-8; **Fig. 1J**), supporting a requirement for endothelial *Piezo1* beyond development to maintain IOP homeostasis in adulthood.

### PIEZO1 activation promotes ANGPT2 secretion and triggers downstream TIE2 signaling in SC

Having established the importance of SC endothelial PIEZO1 in IOP homeostasis and AHO, we next set out to identify the molecular mechanisms underlying this functional role and to determine whether PIEZO1 in SC endothelial cells forms a functional signaling axis with ANGPT2, as it does in lymphatic vessels^17^. To assess whether PIEZO1 signals through its Ca^2+^ channel activity in SC endothelium ex vivo, we generated Cdh5-CreERT2; Salsa6f mice for endothelial-specific Ca^2+^ sensing and induced recombination with tamoxifen. Eyes were dissected, a limbal strip containing SC was mounted, and either vehicle (PBS + 0.5% DMSO) or the PIEZO1 activator Yoda1 (20 µM) was applied. Time-lapse imaging of the green/tdTomato (G/R) ratio in SC endothelial cells showed a greater increase with Yoda1, indicating PIEZO1-mediated Ca^2+^ influx (**Fig. 2A**; n = 4 strips per group). Next, to determine whether PIEZO1 activation induces ANGPT2 release and downstream TIE2 signaling in vivo, we activated PIEZO1 channels in SC endothelium using the small-molecule agonist Yoda1. Eyes received intracameral injection of vehicle (PBS containing 0.5% DMSO) or Yoda1 (20 µM in vehicle, 1 µL) at 100 nL/min; 30 min later, eyes were enucleated, SC whole mounts were prepared, and samples were imaged by confocal microscopy. Vehicle-treated and Yoda1-treated eyes were analyzed as independent groups. Yoda1 treatment led to reduced intracellular ANGPT2 staining, consistent with secretion, together with a concomitant increase in phosphorylated TIE2 (p-TIE2) within the same SC regions (**Fig. 2B**; ANGPT2 intensity, vehicle: 8.788 ± 0.840 vs Yoda1: 5.272 ± 0.614 AU, n = 8 eyes per group, P = 0.0045; p-TIE2 intensity, vehicle: 9.210 ± 2.463 vs Yoda1: 16.354 ± 2.050 AU, n = 5-7 eyes per group, P = 0.0494; unpaired two-tailed Student’s t-test). This acute ANGPT2 release and associated TIE2 phosphorylation was accompanied by activation of downstream signaling, as indicated by increased p-AKT (**Fig. 2C**; unpaired two-tailed t-test, n = 8–13 eyes per group, P = 0.0401) and nuclear export of FOXO1, reflected by a decrease in the FOXO1 nuclear/total intensity ratio in SC endothelial cells (vehicle: 1.214 ± 0.044 vs Yoda1: 1.059 ± 0.011; unpaired two-tailed t-test, n = 3–5 eyes per group, P = 0.022; **Fig. 2D**).

**Figure 2.**
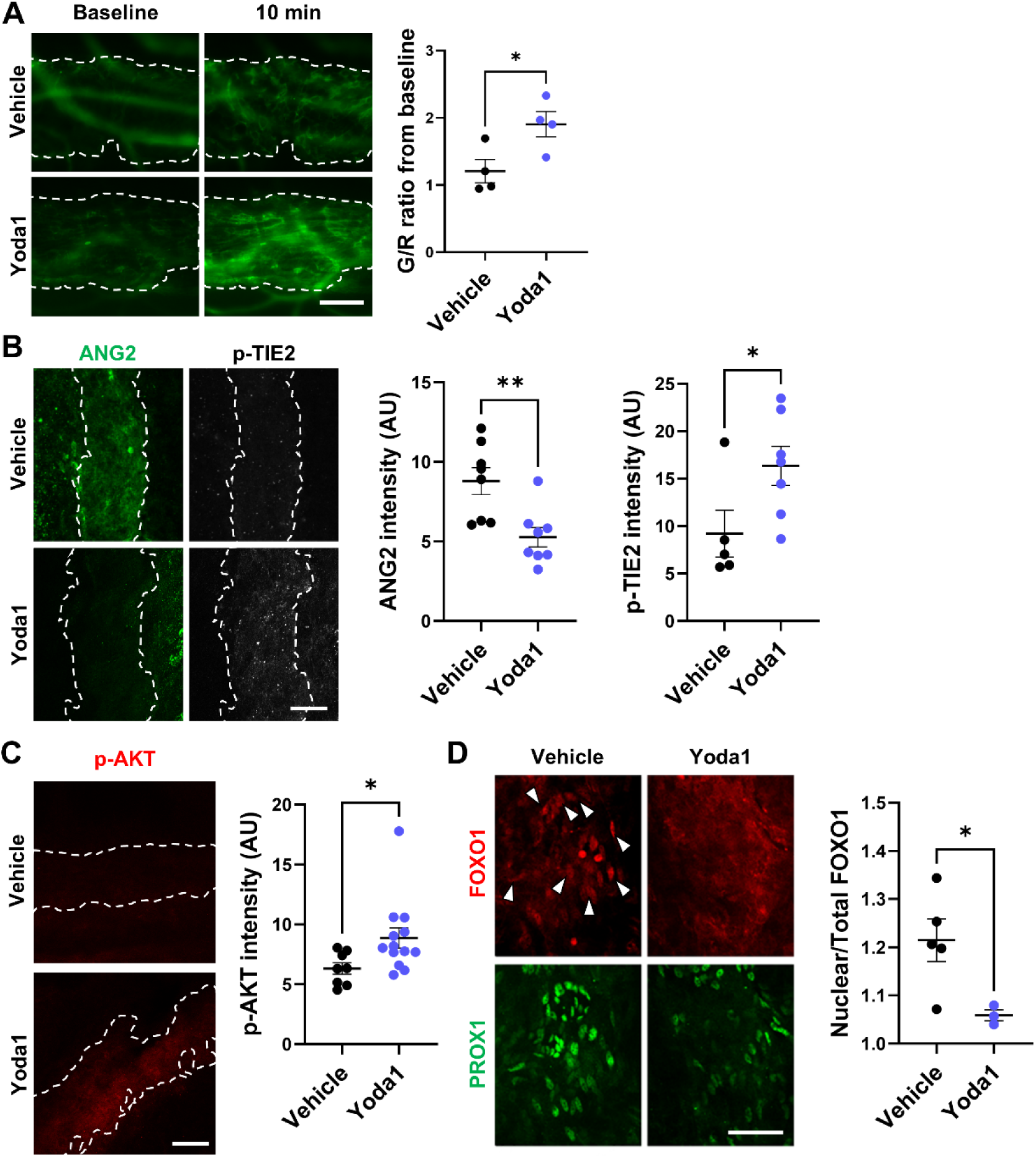
PIEZO1 activation in Schlemm’s canal (SC) induces Ca^2+^ influx and acutely mobilizes ANGPT2 to engage downstream TIE2–AKT–FOXO1 signaling. (A) Ex vivo ratiometric Ca^2+^ imaging of SC endothelium in limbal strips from Cdh5-CreERT2; Salsa6f mice treated with vehicle (PBS + 0.5% DMSO) or Yoda1 (20 µM). Representative images at baseline and 10 min; dashed lines outline SC. Right, change in GCaMP6f/tdTomato (G/R) ratio from baseline. (B) ANGPT2 and phosphorylated TIE2 (p-TIE2) immunostaining on limbal SC whole-mounts 30 min after intracameral injection of vehicle or Yoda1 (20 µM in vehicle, 1 µL at 100 nL/min). Left, ANGPT2 (green); right, p-TIE2 (white/greyscale); dashed lines outline SC. Yoda1 reduces intracellular ANGPT2 and concomitantly increases p-TIE2 in the same regions. Right, quantification of ANGPT2 and p-TIE2 intensities. (C) p-AKT immunostaining (red) is elevated in Yoda1-treated SC. Right, quantification of p-AKT intensity. (D) Co-immunostaining of FOXO1 (red) and PROX1 (green). Arrowheads indicate PROX1^+^ nuclei with prominent nuclear FOXO1 signal in vehicle-treated SC endothelium, which is reduced after Yoda1. Right, quantification of the FOXO1 nuclear-to-total intensity ratio in PROX1^+^ cells. All images were acquired with identical settings. Quantifications were analyzed using an unpaired two-tailed Student’s t-test. Scale bars: (A–C) 100 µm; (D) 50 µm.

### PIEZO1-induced ANGPT2 signaling promotes junctional clustering and activation of integrin α9β1

We have previously reported that Yoda1 treatment increased ITGA9 expression in cultured human lymphatic endothelial cells, suggesting a link between *Piezo1*-mediated Ca^2+^ influx and integrin signaling^17^. In SC, in addition to Ca^2+^ influx and ANGPT2 secretion, we observed increased total ITGA9 immunoreactivity in SC endothelium following intracameral Yoda1 treatment and observed that ITGA9 localization was more linear, with a junction-associated staining pattern (**Supplementary Fig. S1**). Quantification confirmed increased ITGA9 intensity in SC after Yoda1 (vehicle, 5.44 ± 0.42 AU; Yoda1, 7.47 ± 0.66 AU; n = 11–12 eyes; P = 0.0179; **Supplementary Fig. S1**).

Because this antibody does not specifically recognize an active conformation of ITGA9, this result was encouraging, though we interpreted it as enrichment/redistribution rather than direct evidence of ITGA9 activation. In addition to its established role as a TIE2 ligand, ANGPT2 regulates angiogenesis and endothelial cell behavior through integrin engagement, independent of TIE2^16,28,29^. To test whether ANGPT2 is required for PIEZO1-driven junctional enrichment of integrin α9β1, LECs in vitro were transfected with siControl or si*ANGPT2* and then treated with vehicle or Yoda1 prior to fixation and staining. Consistent with our in vivo observations in mouse SC, Yoda1 increased junctional ITGA9 enrichment, reflected by increased ZO-1/ITGA9 colocalization in siControl cells (Pearson’s r: siControl + vehicle, - 0.0608 ± 0.0164; siControl + Yoda1, 0.0567 ± 0.0221; P = 0.0074; **Fig. 3A**). In contrast, ANGPT2 knockdown blunted this Yoda1-induced increase (Pearson’s r: si*ANGPT2* + vehicle, -0.0464 ± 0.0361; si*ANGPT2* + Yoda1, -0.0520 ± 0.0191; P = 0.9986), and ZO-1/ITGA9 colocalization was lower in si*ANGPT2* + Yoda1 compared with siControl + Yoda1 (P = 0.0050; **Fig. 3A**). Because ITGA9 functions as a heterodimer with integrin β1 (ITGB1), we next assessed ITGB1 localization under the same conditions. Yoda1 similarly increased ZO-1/ITGB1 colocalization in siControl cells (Pearson’s r: siControl + vehicle, 0.0383 ± 0.0264; siControl + Yoda1, 0.2059 ± 0.0256; P = 0.000334; **Fig. 3B**). In contrast, this Yoda1-induced increase was not observed following ANGPT2 knockdown (Pearson’s r: si*ANGPT2* + vehicle, 0.1035 ± 0.0381; si*ANGPT2* + Yoda1, 0.1296 ± 0.0220; P = 0.9245; **Fig. 3B**). We next asked whether ANGPT2 was required for PIEZO1-driven integrin activation in SC in vivo.

**Figure 3.**
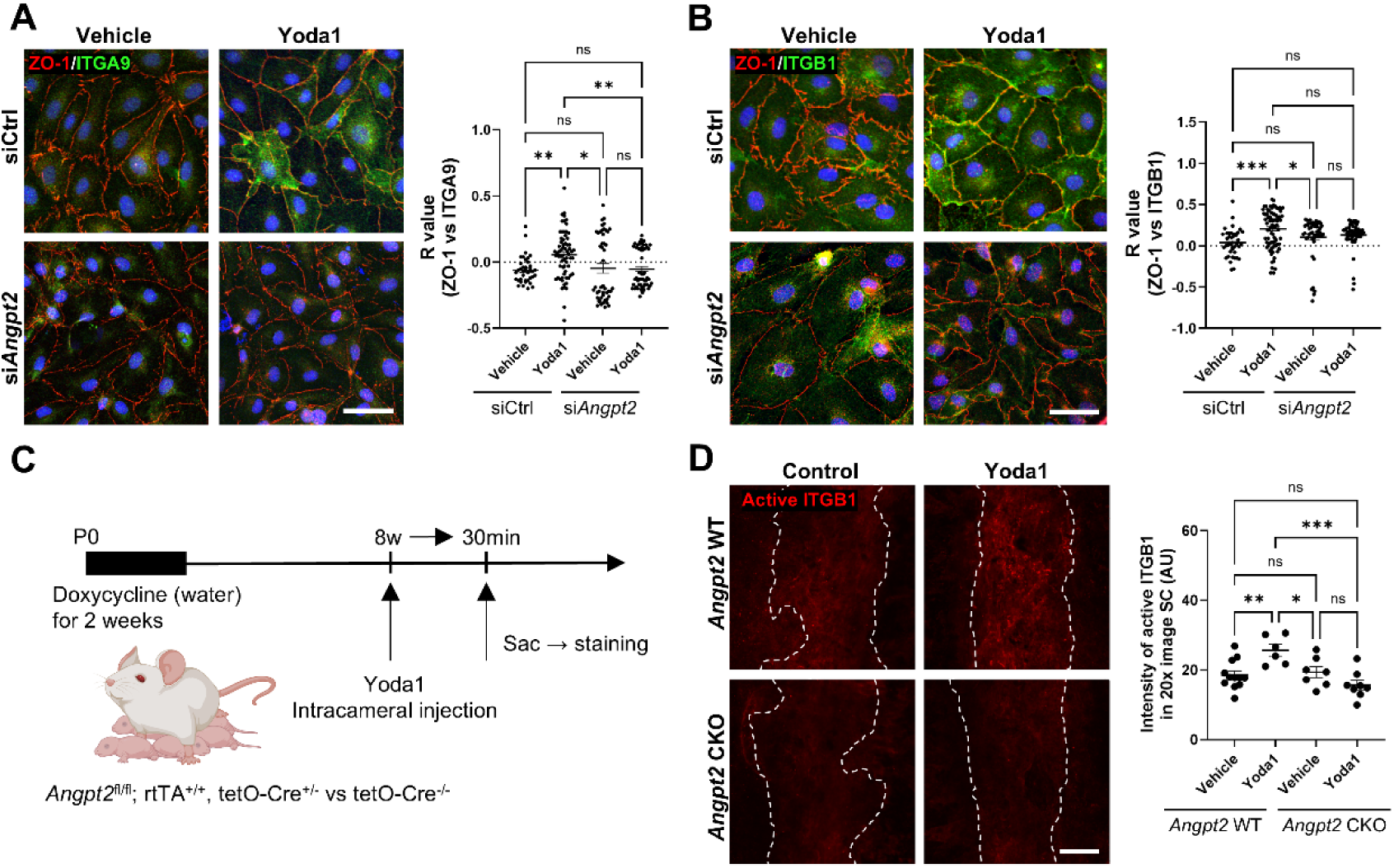
PIEZO1-induced ANGPT2 signaling promotes junctional clustering of integrin α9β1 in vitro and activates β1 integrin in SC. (A) HDLECs transfected with siCtrl or si*Angpt2* and treated with vehicle or Yoda1 (0.5 µM, 5 min) were stained for ZO-1 (red) and ITGA9 (green); Yoda1 increased junctional ZO-1/ITGA9 colocalization in siCtrl cells, which was blunted by *Angpt2* knockdown; Pearson’s correlation coefficient (r) was calculated using Fiji (Coloc2). (B) Under the same conditions, ZO-1 (red) and ITGB1 (green) colocalization increased with Yoda1 in siCtrl cells and was reduced by *Angpt2* knockdown. (C) Experimental scheme for inducible *Angpt2* CKO mice (*Angpt2*^fl/fl^; rtTA^+/+^; tetO-Cre^+/−^ vs tetO-Cre^−/−^) with doxycycline administration in drinking water from P0 for 2 weeks and subsequent intracameral injection of vehicle or Yoda1, followed by SC whole-mount staining 30 min after injection. (D) Active ITGB1 immunostaining in SC whole-mounts 30 min after intracameral injection of vehicle or Yoda1 in *Angpt2* WT and *Angpt2* CKO mice; dashed lines outline SC; right, quantification of active ITGB1 intensity in SC per 20× image. Data are presented as mean ± SEM. Statistics: one-way ANOVA with Tukey–Kramer post hoc test. Scale bars: (A, B) 50 µm; (D) 100 µm.

Inducible *Angpt2* CKO mice were generated and recombination was induced by doxycycline administration (**Fig. 3C**). At 8 weeks of age, mice received unilateral intracameral injection of vehicle or Yoda1 (20 μM) and SC was analyzed 30 min later. Using an antibody recognizing the active conformation of integrin β1, Yoda1 increased active ITGB1 immunoreactivity in SC in *Angpt2* WT controls (WT vehicle, 18.60 ± 1.18 AU; WT Yoda1, 25.69 ± 1.69 AU; P = 0.0100), whereas this response was not observed in *Angpt2* CKO mice (CKO vehicle, 18.82 ± 1.49 AU; CKO Yoda1, 15.82 ± 1.61 AU; P = 0.5142; **Fig. 3D**). Consistent with ANGPT2 dependence, active ITGB1 signal in CKO + Yoda1 was lower than WT + Yoda1 (P = 0.0011; **Fig. 3D**). Altogether, these results indicate that PIEZO1-induced ANGPT2 signaling promotes junctional clustering of integrin α9β1 in vitro and is required for PIEZO1-driven ITGB1 activation in SC in vivo.

### PIEZO1-induced ANGPT2 release activates ITGA9–FAK signaling and rapidly remodels adherens junctions to support aqueous humor outflow

Having established that PIEZO1-induced ANGPT2 signaling promotes junctional enrichment and activation of integrin α9β1, we next examined FAK, a canonical downstream effector of integrins. In human dermal lymphatic endothelial cells (HDLECs), Yoda1 increased FAK phosphorylation (p-FAK) without altering total FAK (p-FAK/FAK: vehicle, 1.00 ± 0.03 vs Yoda1, 1.55 ± 0.13; n = 3; P = 0.0299; **Fig. 4A**). This Yoda1-induced p-FAK response was attenuated by ITGA9 knockdown (ITGA9 reduced by 98.4%; **Supplementary Fig. S2A**) (p-FAK/FAK: siControl + Yoda1, 1.55 ± 0.13 vs siITGA9 + Yoda1, 0.98 ± 0.14; n = 3 per group; P = 0.0240; **Fig. 4A**) and by ANGPT2 knockdown (ANGPT2 reduced by 97.0%; **Supplementary Fig. S2B**) (p-FAK/FAK: siControl + Yoda1, 1.39 ± 0.04 vs siANGPT2 + Yoda1, 0.45 ± 0.02; n = 3 per group; P = 2.68×10^-5^; **Fig. 4B**), indicating that PIEZO1-driven FAK phosphorylation requires both ITGA9 and ANGPT2 (**Fig. 4A,B**). Recombinant ANGPT2 alone was sufficient to elevate p-FAK (p-FAK/FAK: vehicle, 1.00 ± 0.08 vs recombinant ANG2 [rANG2], 2.65 ± 0.06; n = 3; P = 8.36×10^-5^; **Fig. 4D**), supporting an extracellular ANGPT2-dependent mechanism. Consistent with this, neutralization of ANGPT2 with the blocking antibody MEDI3617 suppressed Yoda1-induced p-FAK (p-FAK/FAK: Yoda1, 2.13 ± 0.04 vs MEDI3617 + Yoda1, 0.82 ± 0.25; P = 0.0006; **Fig. 4C**). We next tested whether ANGPT2-driven integrin–FAK signaling is TEK-dependent by knocking down TEK in HDLECs. Despite efficient TEK depletion (TEK reduced by 95.2%; **Supplementary Fig. S2C**), Yoda1 still robustly increased p-FAK (p-FAK/FAK: vehicle, 1.00 ± 0.12 vs Yoda1, 2.20 ± 0.10; n = 3; P = 0.00167; **Fig. 4E**), consistent with a TEK-independent route to ANGPT2-driven FAK activation. To determine whether PIEZO1 activation similarly engages FAK signaling in SC in vivo, we delivered Yoda1 intracamerally and quantified p-FAK immunoreactivity in SC whole mounts. Intracameral Yoda1 increased p-FAK signal in SC compared with vehicle controls (vehicle, 4.72 ± 0.14 AU vs Yoda1, 9.09 ± 1.65 AU; n = 9–10 eyes per group; unpaired two-tailed t-test, P = 0.0233; **Fig. 4F**), indicating that PIEZO1 activation acutely enhances FAK signaling within the conventional outflow pathway. Given that integrin–FAK signaling is tightly coupled to actomyosin remodeling and adherens junction dynamics^30,31^, we next asked whether PIEZO1-induced ANGPT2 release drives rapid junctional remodeling. LEC monolayers were treated with vehicle or Yoda1 in the presence of siControl or siANGPT2 and stained for VE-cadherin. Yoda1 induced a clear thinning and streamlining of VE-cadherin–positive junctions, whereas ANGPT2 knockdown blunted this junctional response, consistent with ANGPT2-dependent junctional remodeling downstream of PIEZO1 (**Fig. 4G**). Finally, to test whether acute extracellular ANGPT2 neutralization is sufficient to alter outflow resistance at the tissue level, eyes were mounted on the iPerfusion system and perfused intracamerally with MEDI3617 (anti-ANGPT2, human IgG1κ; 4 mg/mL) or a matched human IgG1κ isotype control (4 mg/mL) throughout the facility measurement. Outflow facility was reduced in MEDI3617-treated eyes (IgG, 6.49 ± 1.43 vs MEDI3617, 2.27 ± 0.53 nL/min/mmHg; n = 8 paired eyes), corresponding to an approximately 65% reduction; paired t-test on ln(C_r_), P = 0.0030 (**Fig. 4H**). Together, these results support a model in which PIEZO1-induced extracellular ANGPT2 signaling engages ITGA9-dependent integrin–FAK pathways, promotes rapid junctional remodeling, and acutely contributes to aqueous humor outflow.

**Figure 4.**
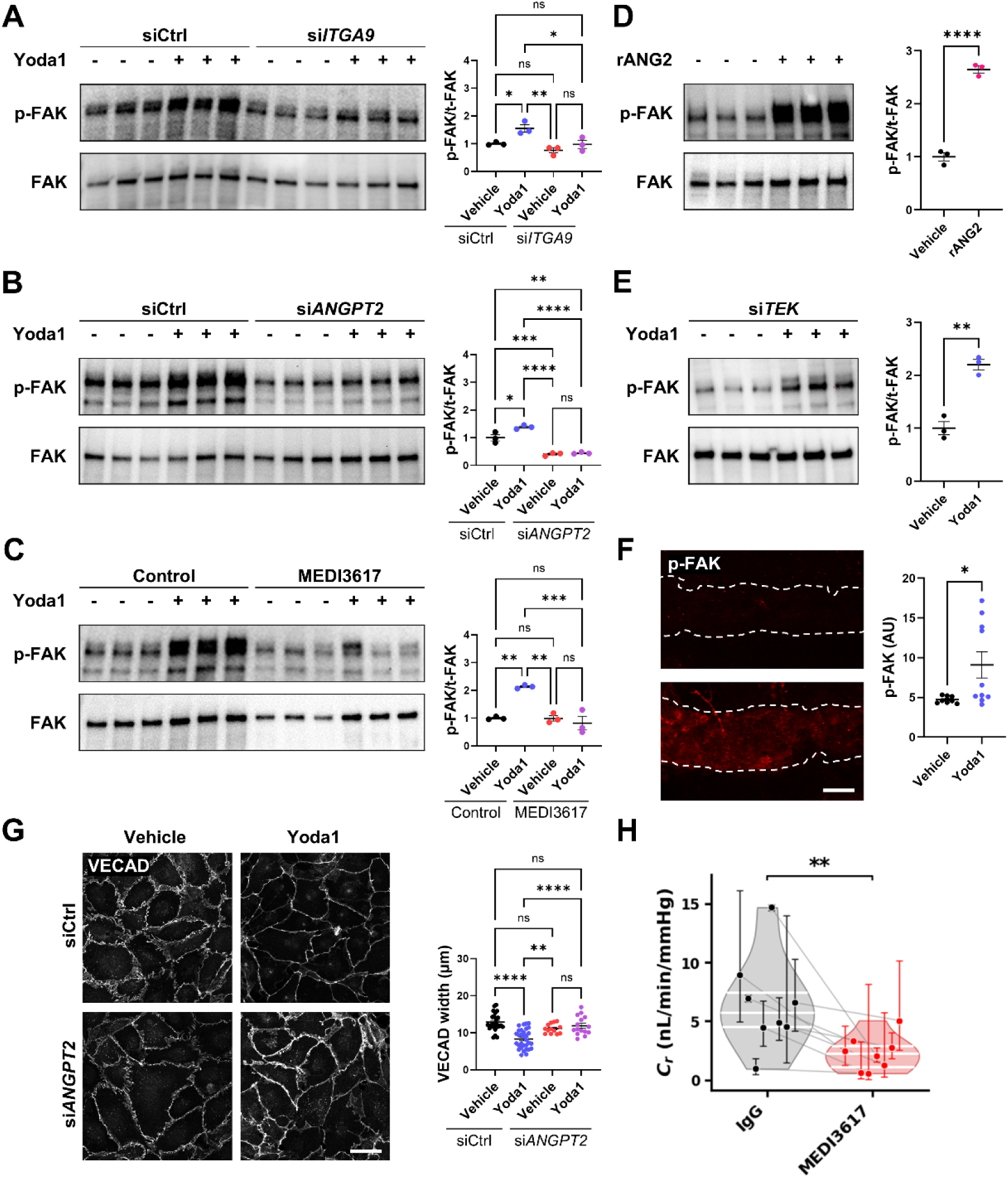
ANGPT2–ITGA9 signaling drives *TEK*-independent FAK phosphorylation downstream of PIEZO1 and is engaged in SC in vivo. (A) Representative immunoblots of p-FAK and total FAK in HDLECs transfected with control siRNA (siCtrl) or si*ITGA9* and treated with vehicle or Yoda1; right, p-FAK/FAK quantification. (B) Representative immunoblots of p-FAK and total FAK in HDLECs transfected with siCtrl or si*ANGPT2* and treated with vehicle or Yoda1; right, p-FAK/FAK quantification showing that *ANGPT2* knockdown attenuates Yoda1-induced p-FAK. (C) ANG2 neutralization (MEDI3617) suppresses Yoda1-induced p-FAK in HDLECs; right, p-FAK/FAK quantification. (D) Recombinant ANG2 (rANG2) is sufficient to increase p-FAK in HDLECs; right, p-FAK/FAK quantification. (E) *TEK* knockdown (si*TEK*) does not prevent Yoda1-mediated FAK phosphorylation; right, p-FAK/FAK quantification. (F) In vivo, p-FAK immunostaining in SC is increased 30 min after intracameral injection of Yoda1 versus vehicle; right, quantification. (G) Yoda1 induces VE-cadherin (VECAD) junctional remodeling, which is blunted by *ANGPT2* knockdown; right, quantification of VECAD width. (H) Ex vivo outflow facility (C_r_) is reduced by continuous intracameral perfusion with MEDI3617 compared with IgG control in paired eyes. Images were acquired with identical settings within each experiment; data are presented as mean ± SEM; Statistics: one-way ANOVA with Tukey–Kramer post hoc test (A–C, G), unpaired two-tailed Student’s t-test (D–F) and paired two-tailed t-test on ln(C_r_) (H). Scale bars: (F) 100 µm; (G) 50 µm.

### Itga9 deficiency impairs SC morphogenesis and outflow function, leading to ocular hypertension and RGC loss

Our single-cell RNA sequencing analysis^26^ revealed that *Itga9* is predominantly expressed in SC endothelial cells, with minimal expression in TM (**Fig. 5A**), consistent with prior reports that SC endothelium expresses integrin α9^8^. While endothelial *Itgb1* (β1 integrin) deletion has been reported to impair SC morphogenesis^32^, the functional contribution of *Itga9* to SC formation or maintenance has not been directly tested. We therefore generated a doxycycline-inducible conditional knockout (*Itga9*^fl/fl^; rtTA^+/+^; tetO-Cre^+/−^; hereafter *Itga9* CKO). Because exons 1–8 are in-frame and exon 9 is also a multiple of three nucleotides, Cre-mediated excision of exon 9 is predicted to preserve the open reading frame and thus produce a truncated ITGA9 protein rather than a complete null allele. Consistent with exon 9 excision, reverse transcription PCR spanning the *Itga9* exon 9 region followed by Sanger sequencing confirmed loss of exon 9 in recombined cDNA (**Supplementary Fig. S3**). Immunoblotting of lung, liver, and kidney lysates demonstrated that full-length ITGA9 was markedly reduced in homozygous recombined tissues compared with heterozygous recombined and non-recombined controls, with an additional lower-molecular-weight immunoreactive band consistent with a residual truncated ITGA9 species (**Supplementary Fig. S3**). Moreover, in lung sections—where the altered pattern is most readily apparent—ITGA9 immunofluorescence in recombined mice was redistributed from a membrane-associated pattern to a predominantly intracellular distribution in endothelial cells, supporting impaired plasma-membrane localization and suggesting loss of

**Figure 5.**
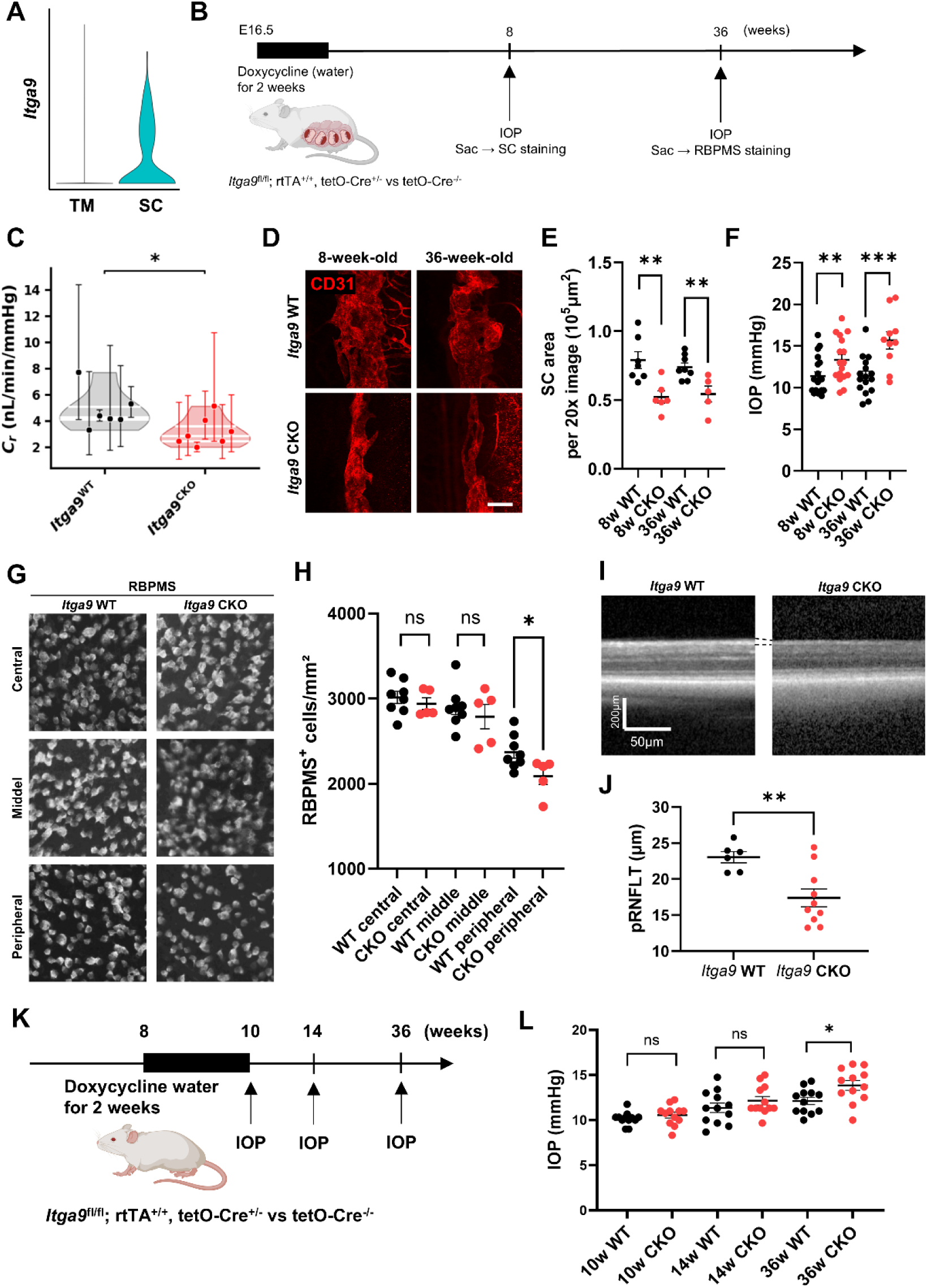
*Itga9* deficiency reduces outflow facility, impairs SC development, elevates IOP, and causes peripheral RGC loss. (A) Violin plot showing *Itga9* expression levels derived from single-cell RNA sequencing of anterior segment tissues, demonstrating selective expression in SC endothelial cells compared with trabecular meshwork (TM). (B) Schematic of the experimental timeline for doxycycline-inducible developmental *Itga9* deletion. Doxycycline was administered from embryonic day 16.5 (E16.5) for 2 weeks to induce Cre expression in *Itga9*^fl/fl^; rtTA^+/+^; tetO-Cre^+/−^ (*Itga9* CKO) mice, with tetO-Cre^−^ littermates as controls, followed by assessment of SC morphometry, outflow facility and IOP at 8 weeks, and RBPMS staining at 36 weeks. (C) Outflow facility (C_r_) is reduced in *Itga9* CKO compared with *Itga9* WT controls at 8 weeks. (D) Representative SC whole-mount images immunostained for CD31 to delineate SC morphology at 8 weeks and 36 weeks. (E) Quantification of SC area shows a significant reduction in *Itga9* CKO mice at both 8 weeks and 36 weeks compared with age-matched *Itga9* WT littermates. (F) IOP measured by rebound tonometry is elevated in *Itga9* CKO mice at both 8 and 36 weeks of age. (G) Representative retinal flat mounts immunostained for RBPMS to label RGCs from *Itga9* WT and *Itga9* CKO mice at 36 weeks, shown for central, middle, and peripheral regions. (H) Quantification of RBPMS-positive RGCs reveals a significant decrease in RGC density in the peripheral retina of *Itga9* CKO mice, with no difference in the central or middle regions. (I) Representative peripapillary OCT B-scan centered on the optic nerve head (ONH); RNFL boundaries are indicated. (J) Quantification of peripapillary retinal nerve fiber layer thickness (pRNFLT) between *Itga9* WT and *Itga9* CKO. (K) Schematic of the experimental timeline for postnatal *Itga9* deletion using doxycycline-inducible tetO-Cre. Doxycycline was administered from 8 to 10 weeks of age in *Itga9*^fl/fl^; rtTA^+/+^; tetO-Cre^+/−^ (*Itga9* CKO) mice, with Cre^−^ littermates as controls. IOP was measured at 10, 14, and 36 weeks. (L) Longitudinal IOP measurements show no difference at 10 or 14 weeks, but significantly elevated IOP in *Itga9* CKO mice at 36 weeks. Statistical analyses were performed using two-tailed unpaired Student’s t-test for two-group comparisons, with comparisons performed within each age/region/time point as indicated in the panels. Data are presented as mean ± SEM. *P < 0.05, **P < 0.01, ***P < 0.001, ns = not significant. Scale bars: (D) 100 µm; (G) 50 µm; (I) 200 µm (vertical), 50 µm (horizontal).

ITGA9 function despite residual protein expression (**Supplementary Fig. S3**). Doxycycline was administered from embryonic day 16.5 (E16.5) for two weeks to induce *Itga9* deletion during fetal development (**Fig. 5B**). Cre-negative littermates (*Itga9*^fl/fl^; rtTA^+/+^; tetO-Cre^−^) served as controls. Outflow facility was reduced in *Itga9* CKO compared with controls (WT, 4.83 ± 1.54 vs CKO, 3.16 ± 1.08 nL/min/mmHg; n = 6–7 eyes per group; P = 0.028; **Fig. 5C**). CD31 immunostaining at 8 weeks and 36 weeks revealed a significant reduction in SC area in *Itga9* CKO at both ages (**Fig. 5D,E**; unpaired two-tailed t-tests within age, n = 5–8 eyes per group; 8 w: P < 0.01; 36 w: P < 0.01). Consistently, IOP was higher in CKO than controls at 8 weeks and 36 weeks (**Fig. 5F**; unpaired two-tailed t-tests within age, n = 10–19 eyes per group; 8 w: P < 0.01; 36 w: P < 0.001), indicating that *Itga9* is required for SC development and IOP homeostasis. To assess the consequences of chronic IOP elevation, we quantified RBPMS-positive RGC density at 36 weeks on retinal flat mounts and compared genotypes within each retinal eccentricity, and we measured peripapillary retinal nerve fiber layer thickness (pRNFLT) by optical coherence tomography (OCT). RGC density was significantly decreased in the peripheral retina of *Itga9* CKO, with no difference in central or middle regions (**Fig. 5G,H**; n = 5–8 eyes per group; peripheral: P < 0.05; central/middle: ns), suggesting region-specific RGC loss and phenocopying *Piezo1*^ΔEC^. In parallel, pRNFLT was reduced in *Itga9* CKO compared with WT (**Fig. 5I,J**; n = 6–10 mice; P < 0.01). Finally, to test whether *Itga9* is required for adult IOP homeostasis, we induced postnatal deletion by administering doxycycline in drinking water from 8 to 10 weeks of age in *Itga9*^fl/fl^; rtTA^+/+^; tetO-Cre^+/−^ mice (*Itga9* CKO), with Cre^−^ littermates as controls (**Fig. 5K**). IOP did not differ between genotypes at 10 or 14 weeks (10 w: WT, 10.17 ± 0.21 vs CKO, 10.56 ± 0.32 mmHg, P = 0.319; 14 w: WT, 11.36 ± 0.52 vs CKO, 12.16 ± 0.47 mmHg, P = 0.266; n = 12 mice per group), but was significantly elevated at 36 weeks in *Itga9* CKO (WT, 12.12 ± 0.40 vs CKO, 13.85 ± 0.55 mmHg; P = 0.0186; n = 12 mice per group; **Fig. 5L**), indicating that *Itga9* is required not only during development but also for long-term IOP homeostasis in adulthood. Consistent with reduced outflow facility in *Itga9* CKO eyes, qualitative transmission electron microscopy (TEM) examination of the SC inner wall revealed readily apparent giant vacuoles (GVs) in WT eyes, whereas GVs were less prominent in *Itga9* CKO eyes under the same processing conditions (**Supplementary Fig. S4**). This finding is consistent with the lower mean number of GVs per 100 μm of SC inner wall in *Itga9* CKO eyes than in WT eyes (3.17 ± 0.93 vs. 8.84 ± 2.26).

### Heterozygous deletion of *Piezo1* and *Itga9* reveals combined effects on SC narrowing and IOP elevation

We showed above that deleting *Piezo1* or *Itga9* alone reduces SC area and elevates IOP, yielding similar glaucomatous phenotypes. These observations suggested that the two genes act along a common axis in SC in vivo, and that partial loss of both might produce a combined effect. To test this, we created germline-deleted heterozygous alleles (**Supplementary Fig. S5A**): *Piezo1*^fl/fl^ or *Itga9*^fl/fl^ mice were crossed with EIIa-Cre to induce germline deletion. The resulting *Piezo1*^+/−^ or *Itga9*^+/−^ offspring were then crossed with WT mice to remove EIIa-Cre. Finally, *Piezo1*^+/−^ and *Itga9*^+/−^ carriers were intercrossed to generate four genotypes: *Itga9*^+/+^; *Piezo1*^+/+^, *Itga9*^+/−^; *Piezo1*^+/+^, *Itga9*^+/+^; *Piezo1*^+/−^, and *Itga9*^+/−^; *Piezo1*^+/−^. At 8 weeks of age, IOP differed across genotypes (**Supplementary Fig. S5C**; one-way ANOVA with Tukey’s multiple comparisons, n = 8–15 mice per genotype): both single heterozygotes were higher than WT (Tukey-adjusted P = 0.0206 for *Itga9*^+/−^; *Piezo1*^+/+^ and P = 0.0052 for *Itga9*^+/+^; *Piezo1*^+/−^), and the double heterozygote was further elevated relative to each single heterozygote (P < 0.0001 vs *Itga9*^+/−^; *Piezo1*^+/+^; P = 0.0121 vs *Itga9*^+/+^; *Piezo1*^+/−^) and to WT (P < 0.0001). For SC area, *Itga9*^+/−^; *Piezo1*^+/+^ was reduced versus WT (P = 0.0042), *Itga9*^+/+^; *Piezo1*^+/−^ showed a trend (P = 0.0691), and the double heterozygote showed the greatest reduction (**Supplementary Fig. S5B,D**; one-way ANOVA with Tukey, n = 4–6), significant versus WT (P < 0.0001) and versus *Itga9*^+/+^; *Piezo1*^+/−^ (P = 0.0246), with a trend versus *Itga9*^+/−^; *Piezo1*^+/+^ (P = 0.0896). Collectively, partial loss of both *Piezo1* and *Itga9* produces a combined effect—most evident for IOP—and drives the largest SC area reduction, supporting functional convergence of these genes in maintaining SC homeostasis and restraining IOP in vivo.

### *ITGA9* knockdown in LECs downregulates flow- and proliferation-related pathways

Although we observed acute changes along the PIEZO1–ANGPT2–integrin α9β1–VE-cadherin axis, these acute effects alone did not fully account for the in vivo phenotype, which included reduced SC area and long-term effects on IOP following adult Itga9 deletion. We therefore next sought to more comprehensively define the downstream events regulated by ITGA9. To this end, we leveraged lymphatic endothelial cells (LECs), which share key molecular features with SC endothelium, and profiled transcriptional changes induced by *ITGA9* knockdown (KD). Total RNA was isolated from control and *ITGA9* KD LECs and subjected to bulk RNA sequencing (bulk RNA-seq). Using genes that were downregulated upon *ITGA9* KD, we performed Kyoto Encyclopedia of Genes and Genomes (KEGG) pathway enrichment analysis, which revealed reduced signatures in pathways consistent with our mechanistic model, including integrin signaling, regulation of actin cytoskeleton, focal adhesion, gap junction, and PI3K–Akt signaling. In addition, we observed enrichment of downregulated genes within KEGG pathways 05418 and 04110, which encompass genes implicated in shear stress sensing and cell cycle, respectively (**Fig. 6A**). To visualize gene-level changes underlying these enrichments, we plotted a heatmap of representative genes from these pathways, revealing coordinated downregulation of core cell-cycle regulators (e.g., *CCND1*/*CCND2*, *CDK1*/*WEE1*) and shear stress–responsive endothelial genes (e.g., *NOS3*, *ICAM1*/*VCAM1*, *TRPV4*) in *ITGA9* KD LECs (**Fig. 6B**).

**Figure 6.**
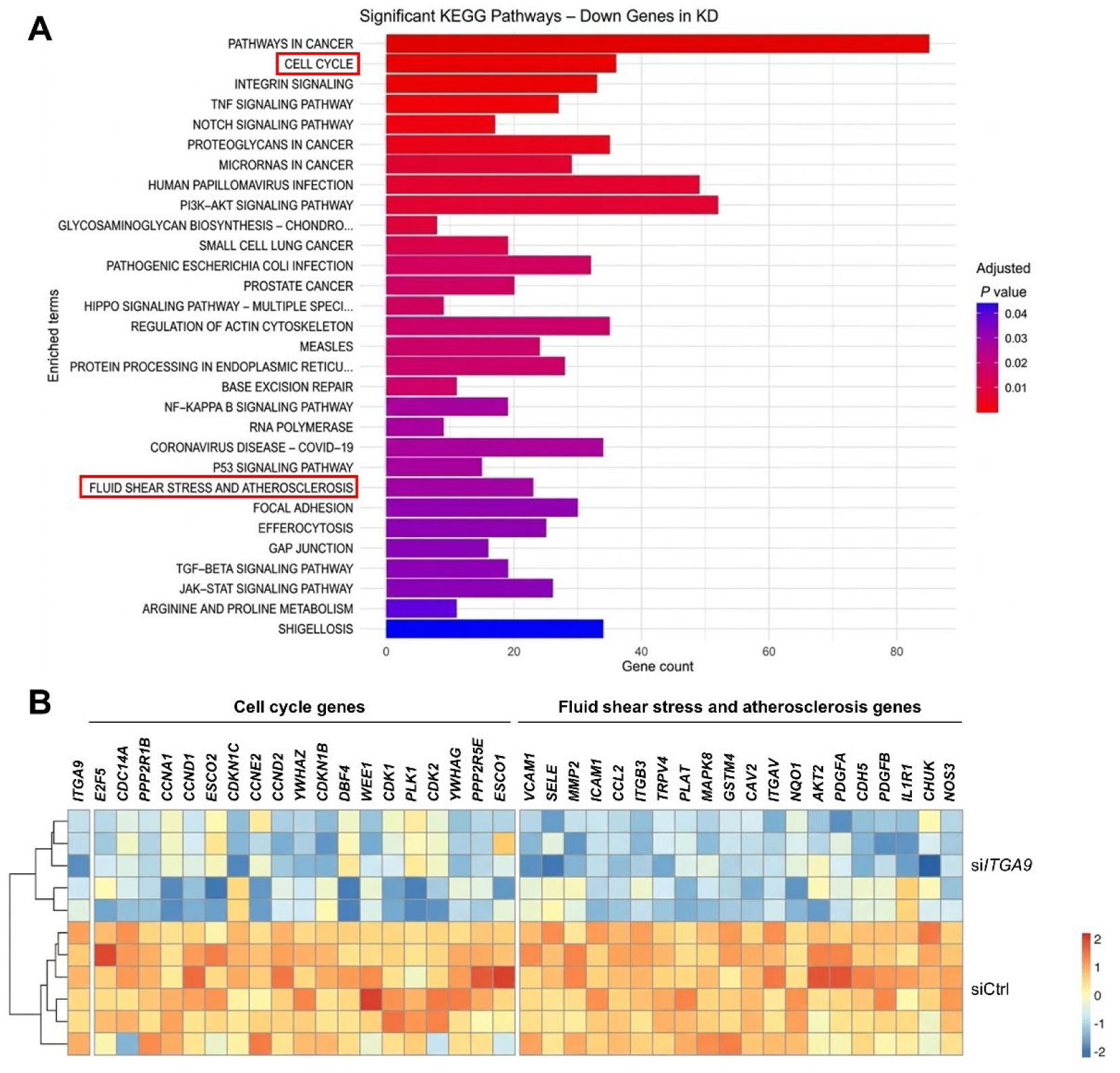
*ITGA9* knockdown downregulates flow- and proliferation-related pathways in lymphatic endothelial cells. (A) Bulk RNA-seq was performed on LECs transfected with siCtrl or si*ITGA9*, and KEGG pathway enrichment analysis was conducted using genes downregulated after *ITGA9* knockdown. Bar length indicates the number of downregulated genes assigned to each enriched KEGG pathway (gene count), and color denotes the adjusted P value. The Cell cycle and Fluid shear stress and atherosclerosis pathways are highlighted. (B) Heatmap showing z-scored expression (per gene) of selected genes annotated to the KEGG Cell cycle and KEGG Fluid shear stress and atherosclerosis pathways, with *ITGA9* included as a knockdown-validation gene, across individual siCtrl and si*ITGA9* samples. Rows represent samples and columns represent genes; hierarchical clustering of samples is shown on the left.

These gene programs are established components of endothelial proliferative and flow-responsive signaling^33–35^. Together, these findings suggest that *ITGA9* contributes to maintaining endothelial proliferative programs and flow-responsive molecular signaling, consistent with the altered flow-dependent proliferative response observed in SC endothelium described below.

### The *Piezo1*–*Itga9* axis promotes outflow-dependent proliferation of SC endothelial cells

PIEZO1 is a mechanosensitive ion channel and is expected to function as a key mechano-sensor in SC, where AHO provides the dominant mechanical input. In lymphatic endothelium, flow/shear activates PIEZO1 to drive endothelial expansion and proliferation^18^. In addition, *Itga9* (integrin α9β1) is essential for lymphatic valve formation and maintenance at flow-defined sites^36^ and regulates endothelial proliferation and migration through extracellular matrix (ECM) coupling^37^. Collectively, these studies suggest *Itga9* participates in angiogenic/organogenetic programs and could contribute to SC development and maintenance. We therefore hypothesized that mechanical stress generated by AHO activates the *Piezo1*–*Itga9* axis to promote SC endothelial proliferation; conversely, loss of *Piezo1* or *Itga9* would blunt flow-induced proliferation, reduce endothelial cellularity, and narrow SC—thereby further reducing flow in a pathogenic feedback loop. AHO in SC is segmental, producing local high- and low-flow regions^8^. To examine the relationship between flow and endothelial proliferation, we injected FluoSpheres™ into the anterior chamber at 3 months of age and, on the same day, administered EdU (5-ethynyl-2′-deoxyuridine) in drinking water (0.2 mg/mL) continuously for 7 days before tissue harvest.

Whole-mount anterior segments were prepared, and for each quadrant we quantified EdU^+^ETS-related gene (ERG)^+^ SC endothelial cells alongside integrated FluoSpheres intensity. FluoSpheres intensity and EdU^+^ERG^+^ counts were positively correlated (Pearson r = 0.643, P = 0.0022), supporting a shear-dependent increase in local SC endothelial proliferation (**Fig. 7A,B**). We next tested genetic requirements. In *Piezo1*^ΔEC^ mice, both the number of SC endothelial cells and the fraction of ERG^+^EdU^+^ cells in the canal were reduced, indicating impaired endothelial proliferation (**Fig. 7C,E–G**). Similarly, doxycycline-inducible *Itga9* CKO (*Itga9*^fl/fl^; rtTA^+/+^; tetO-Cre^+/−^) showed decreased EdU labeling and reduced SC endothelial cellularity (**Fig. 7D,H–J**), phenocopying *Piezo1*^ΔEC^. Together, these results indicate that mechanical cues from AHO are sensed by PIEZO1 and transduced through ITGA9 to drive local SC endothelial proliferation and preserve canal structure in addition to acute VE-cadherin remodeling (**Fig. 8**). Disruption of this axis by loss of *Piezo1* or *Itga9* impairs the proliferative response, leading to reduced endothelial cellularity and canal narrowing, which in turn exacerbates outflow reduction in a deleterious feedback loop.

**Figure 7.**
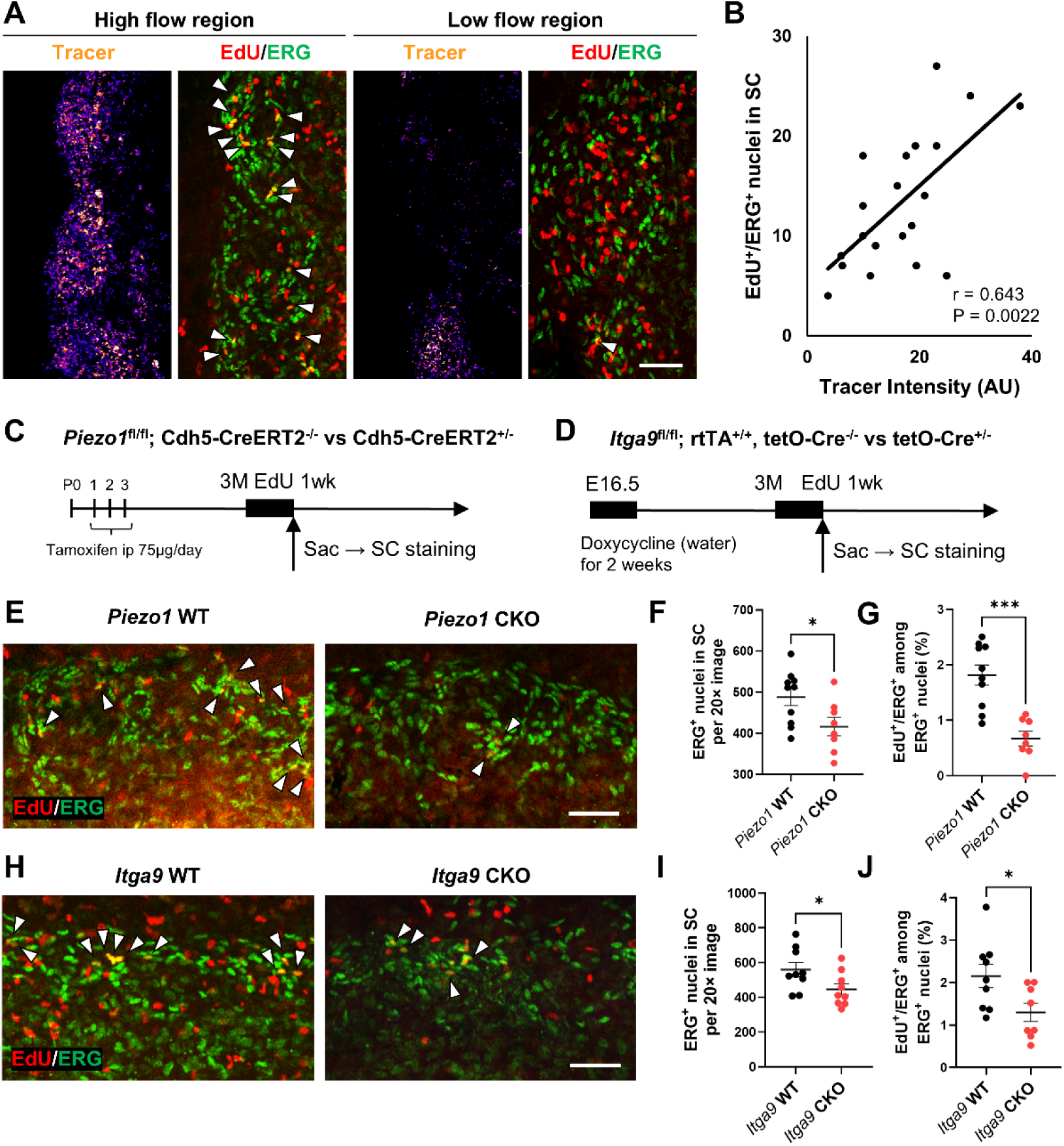
Outflow-dependent proliferation of Schlemm’s canal (SC) endothelium requires the *Piezo1*–*Itga9* axis. (A) SC whole-mounts after intracameral injection of tracer (FluoSpheres™) and 7-day EdU labeling: a high-tracer (high-flow) region shows more EdU^+^/ERG^+^ nuclei in SC than a low-tracer (low-flow) region; tracer (fire), ERG (green), EdU (red); arrowheads mark EdU^+^/ERG^+^ cells. (B) Integrated tracer intensity correlates with the number of EdU^+^/ERG^+^ nuclei in SC (Pearson r = 0.643, P = 0.0022). (C) Experimental scheme for endothelial *Piezo1* deletion: *Piezo1*^fl/fl^; Cdh5-CreERT2^−/−^ vs Cdh5-CreERT2^+/−^; tamoxifen at P1–P3; tracer injection and EdU for 1 week at 3 months; harvest for SC staining. (D) Experimental scheme for inducible *Itga9* deletion: *Itga9*^fl/fl^; rtTA^+/+^; tetO-Cre^−/−^ vs tetO-Cre^+/−^; doxycycline from E16.5 for 2 weeks; tracer injection and EdU for 1 week at 3 months; harvest for SC staining. (E) Representative SC fields from *Piezo1* WT and CKO; arrowheads indicate EdU^+^/ERG^+^ nuclei. (F) Quantification: number of ERG^+^ nuclei in SC per 20× image is reduced in *Piezo1* CKO. (G) Quantification: EdU^+^/ERG^+^ among ERG^+^ nuclei (%) is decreased in *Piezo1* CKO. (H) Representative SC fields from *Itga9* WT and CKO. (I) Quantification: number of ERG^+^ nuclei in SC per 20× image is reduced in *Itga9* CKO. (J) Quantification: EdU^+^/ERG^+^ among ERG^+^ nuclei (%) is decreased in *Itga9* CKO. Data are presented as mean ± SEM. Statistics: (B) Pearson correlation; two-group comparisons (F, G, I, J) used unpaired two-tailed Student’s t-test. *P < 0.05, ***P < 0.001, ns = not significant. Scale bars: (A, E, H) 100 µm.

**Figure 8.**
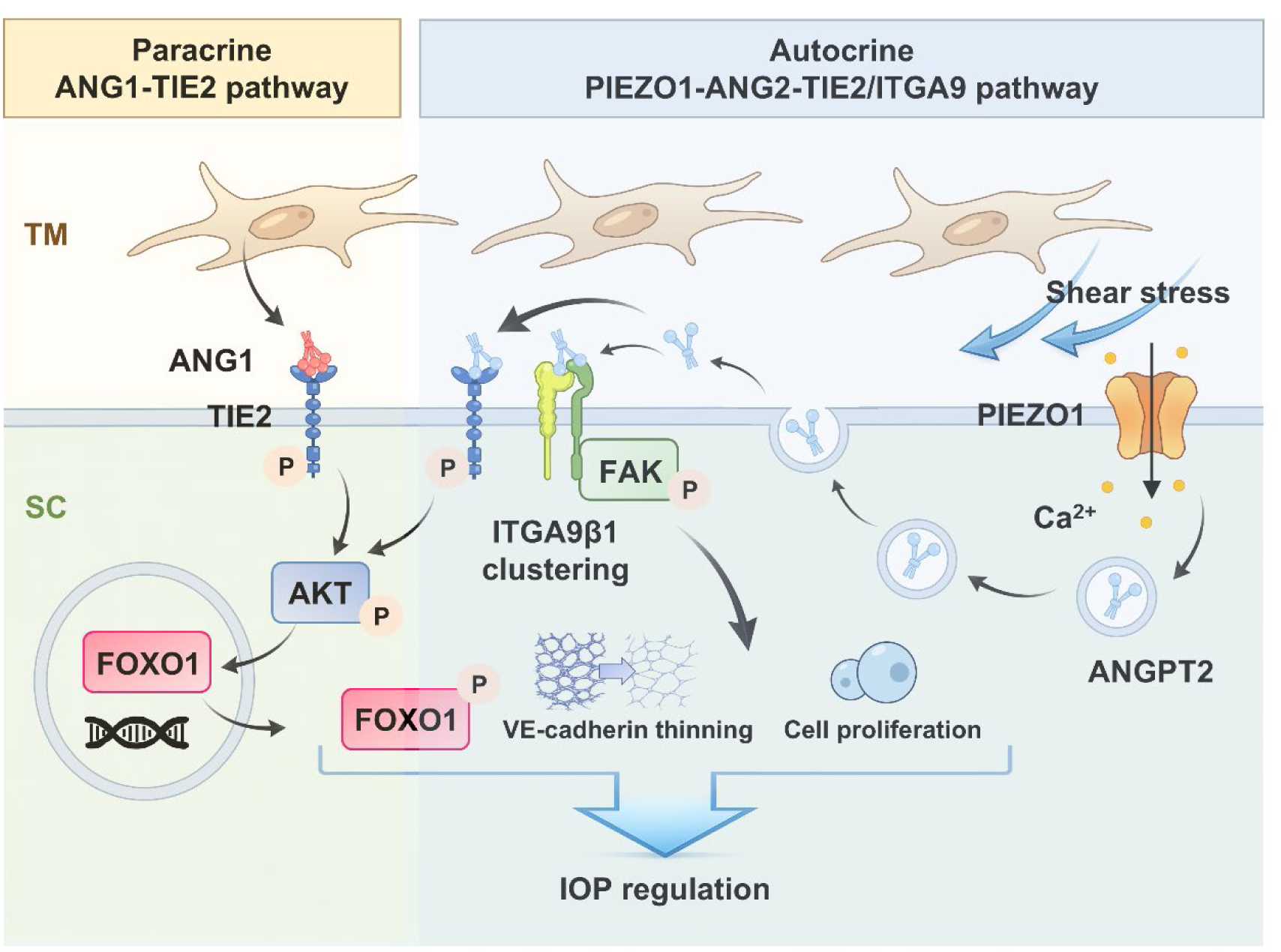
Model of mechanosensitive autocrine PIEZO1–ANGPT2–TIE2/ITGA9 signaling in SC regulating IOP. Schematic summarizing two ANGPT-dependent signaling arms in the conventional outflow pathway. Left, the paracrine ANGPT1–TIE2 pathway: trabecular meshwork (TM)-derived ANGPT1 activates TIE2 in Schlemm’s canal (SC) endothelium, leading to AKT activation and downstream FOXO1 regulation. Right, the autocrine PIEZO1–ANGPT2–TIE2/ITGA9 pathway: shear stress from aqueous outflow activates PIEZO1 in SC endothelium to drive Ca^2+^ influx and ANGPT2 release. Extracellular ANGPT2 engages two downstream arms in SC, including TIE2 phosphorylation/AKT signaling and a TIE2-independent integrin arm characterized by ITGA9β1 clustering and FAK phosphorylation. The integrin–FAK branch promotes rapid adherens junction remodeling (VE-cadherin thinning) and supports flow-dependent SC endothelial proliferation. Together, these paracrine and autocrine pathways converge to maintain SC structure and outflow function, thereby regulating intraocular pressure (IOP).

## Discussion

Throughout life, IOP must be maintained within a narrow range to prevent retinal ganglion cell loss, optic nerve damage, and vision loss. Tissues of the conventional aqueous outflow pathway sense pressure and dynamically regulate outflow resistance through the TM and the inner wall of SC, where aqueous humor traverses a specialized endothelial barrier via transcellular transport through giant vacuoles or paracellular openings^38–40^. However, the molecular mechanisms that translate mechanical sensing into outflow regulation remain poorly understood. Here, we identify a novel mechanosensitive, autocrine ANGPT2 signaling pathway within the SC endothelium that regulates this process, linking fluid-induced mechanical cues, including shear stress, to outflow resistance and IOP homeostasis.

### Mechanotransduction and Endothelial Homeostasis

Across vascular beds, endothelial barriers translate mechanical inputs—fluid shear, stretch, and pressure gradients—into coordinated changes in cytoskeletal tension, junctional organization, and permeability that maintain tissue fluid homeostasis. Disruption of this mechanotransduction compromises barrier adaptability and can drive organ-specific disease^41,42^. In the anterior segment of the eye, mechanical cues are critical regulators of the SC endothelium and AHO, modulating canal caliber and filtration area, inner-wall permeability via transcellular and paracellular routes, and vasodilation of distal outflow vessels^10,43–46^. Recent in vitro studies implicate mechanosensitive Ca^2+^ signaling, including TRPV4, in regulation of the transcellular outflow route in human SC cells^43^. However, how Ca^2+^ signaling controls other determinants of AHO resistance and engages outflow-modulating cytoskeletal, adhesion, and junctional programs in vivo has remained unclear. Because giant vacuoles are pressure-driven membrane deformations associated with high local biomechanical strain in the SC inner wall, their formation and/or the local events that trigger their formation provide a physiologically relevant context for activation of highly tension-sensitive channels. Unlike TRPV4 and other mechanosensitive channels, the PIEZO channels are exquisitely sensitive to lateral stretch, making them plausible candidates to transduce pressure changes generated by AHO to cell behaviors needed to regulate outflow^47,48^.

Here, we show that PIEZO1-dependent Ca^2+^ influx activates an autocrine ANGPT2–integrin α9β1 signaling relay that engages FAK, linking mechanical inputs to SC endothelial remodeling and outflow regulation.

### PIEZO1 as a Mechanosensitive Regulator of SC Endothelium

PIEZO1, a mechanosensitive cation channel, converts membrane tension generated by fluid flow and pressure gradients into intracellular Ca^2+^ signals that shape endothelial cytoskeletal and junctional states^49^. Human genetic studies implicate PIEZO1 variants in primary open-angle glaucoma (POAG) risk and IOP regulation^21^, and pharmacologic studies show that Yoda1 lowers IOP whereas GsMTx4 decreases outflow facility^20,50,51^. Because PIEZO1 is expressed in both the TM and the SC endothelium, however, these studies could not identify the responsible tissue. Here, cell-type–specific deletions show that endothelial Piezo1 loss, rather than TM loss, elevates IOP and reduces outflow facility in mice, supporting the SC endothelium as the primary site of steady-state PIEZO1-mediated outflow regulation. While we cannot exclude transient or other regulatory roles for PIEZO1 within the TM, these findings support a model in which PIEZO1 acts across the conventional outflow pathway, with dominant regulation residing in the SC endothelium.

### ANGPT2 Modulates Integrin Signaling in SC downstream of PIEZO1

Prior studies have shown that PIEZO1 activation triggers rapid ANGPT2 secretion from Weibel–Palade bodies in lymphatic endothelial cells (LECs)^17^. Consistent with a similar mechanism in SC, intracellular ANGPT2 immunostaining in SC endothelial cells rapidly diminished after Yoda1 administration, coincident with increased TIE2 phosphorylation. These findings indicate that PIEZO1-dependent ANGPT2 release can modulate TIE2 signaling in SC. However, unlike LECs, which largely lack ANGPT1-producing pericytes, SC endothelial cells lie adjacent to a persistent source of ANGPT1 produced by juxtacanalicular cells of the TM^12,24^. Given the high basal TIE2 phosphorylation in SC and prior evidence that *Angpt2* deletion in young mice has minimal effects on IOP or SC morphology through the TIE2 signaling pathway^12^, we hypothesized that ANGPT2 may serve an alternative function. In other vascular beds, including the retina, ANGPT2 can activate integrins in a TIE2-independent manner^16,52^. Here, we found that *Angpt2* knockdown almost completely abrogated PIEZO1-induced integrin clustering in mouse SC and FAK activation in vitro, suggesting that ANGPT2 functions primarily as a regulator of integrin signaling in this context. This ANGPT2-mediated integrin signaling axis was TIE2-independent and was not affected by TIE2 knockdown.

### Integrin α9β1 as a Mechanosensitive Outflow Regulator

Integrins are transmembrane receptors that couple the extracellular matrix to the cytoskeleton and can convert mechanical or pressure cues into biochemical signals that regulate cytoskeletal dynamics, proliferation, and adhesion, in part through FAK signaling^53,54^. Integrin signaling is essential for the development and function of conventional outflow tissues, and although integrin biology has been studied most extensively in the TM, recent work demonstrates that expression of the integrin β1 subunit (ITGB1) is required for SC development^32^. In the adult canal, both transcellular (GV) and paracellular (junctional) outflow routes are modulated by pathways that intersect with FAK signaling, including Src family kinases^44^ and Rho kinase–mediated cytoskeletal regulation^55^.

Although multiple ITGB1-associated α subunits are expressed in SC endothelium, the specific contributions of individual α integrins to SC structure and function have remained less well defined^10,56–58^. By establishing the requirement for *Itga9* in FAK phosphorylation downstream of PIEZO1-dependent ANGPT2 release, we provide a mechanistic framework for how SC cells translate IOP- and shear stress–derived signals into regulation of outflow resistance.

### Connecting Genetic Risk to the Mechanosensing Pathway

The biological relevance of the *ANGPT2*–integrin pathway is reinforced by its convergence with established glaucoma risk factors. In addition to the associations between *PIEZO1* and glaucoma discussed above, variants at the *ANGPT2* locus have also been linked to IOP regulation and POAG risk^24,59^.

Furthermore, the interaction between *Itga9* and one of its ligands, *SVEP1* (Polydom), is notable given that *SVEP1* variants have been reported as genetic modifiers in both primary congenital glaucoma (PCG) and POAG^24,60,61^. Our data suggest that these genetic associations may manifest through impaired flow sensing. Because *SVEP1* can also interact with ANGPT ligands, its role as an *Itga9* ligand raises the possibility of a multiprotein complex that stabilizes the SC wall against the mechanical demands of AHO^61^.

### Cell Proliferation Downstream of PIEZO1–ITGA9 Signaling

The observation that loss of *Itga9* or *Piezo1* selectively impairs SC endothelial cell proliferation in high-flow regions is intriguing. SC endothelial cells must preserve a functional barrier while undergoing continual and often pronounced cell shape changes and turnover. Our results suggest that *Itga9* functions not merely as an extracellular matrix anchor, but as a dynamic regulator of the cytoskeleton, enabling the cytoskeletal and junctional remodeling required for proliferation while maintaining barrier integrity under high shear stress.

## Conclusion

In summary, our study defines a specialized mechanotransduction axis in Schlemm’s canal endothelium in which PIEZO1-mediated Ca^2+^ influx triggers an unconventional, TIE2-independent ANGPT2–integrin signaling pathway. By linking mechanical cues such as membrane stretch and/or fluid shear directly to molecular regulators of both acute outflow facility and long-term structural maintenance through the integrin α9β1 complex, these findings provide a new framework for understanding outflow homeostasis and IOP regulation.

Targeting this mechanosensitive autocrine loop may offer therapeutic opportunities to restore pressure sensing and enhance aqueous humor drainage in glaucoma.

## Online methods

### Study approvals

Animal experiments were approved by the Animal Care and Use Committee at Northwestern University (Evanston, IL, USA) and complied with ARVO guidelines for care and use of vertebrate research subjects in Ophthalmology research.

### Animal generation and husbandry

All mice were housed in the Center for Comparative Medicine at Northwestern University (Chicago, IL, USA) under standard conditions (12-hour light/dark cycle, ambient temperature 21–23 °C, 30–70% relative humidity) with ad libitum access to water and standard chow (Teklad #7912, Envigo, Indianapolis, IN, USA). *Itga9*^fl/fl^ mice were a generous gift from Dr. Livingston Van De Water (Albany Medical College), originally described in Singh et al.^62^. *Piezo1*^fl/fl^ mice (B6.Cg-*Piezo1*tm2.1Apat, JAX stock #029213), Cdh5-CreERT2 mice (Tg(Cdh5-cre/ERT2)1Rha, JAX, MGI:3848982), and Rosa26-rtTA mice (Gt(ROSA)26Sortm1(rtTA,EGFP)Nagy, JAX stock #005670) were obtained from The Jackson Laboratory (Bar Harbor, ME, USA). tetO-Cre transgenic mice

(B6.Cg-Tg(tetO-cre)1Jaw/J, JAX stock #006234) and *Wnt1*-Cre mice (B6.Cg-H2az2 Tg(*Wnt1*-cre)11Rth Tg(*Wnt1*-GAL4)11Rth/J, JAX stock #009107) were also obtained from The Jackson Laboratory. LSL-Salsa6f reporter mice (B6(129S4)-Gt(ROSA)26Sortm1.1(CAG-tdTomato/GCaMP6f)Mdcah/J, JAX stock #031968) were also obtained from The Jackson Laboratory.

*Itga9*^fl/fl^ mice were crossed with Rosa26-rtTA and tetO-Cre transgenic mice to generate doxycycline-inducible *Itga9* CKO mice. Pregnant dams were treated starting at embryonic day 16.5 (E16.5) by giving 0.5% (wt/vol) doxycycline-containing drinking water (with 5% sucrose), which was continued for 2 weeks to induce Cre-mediated recombination during fetal development. For postnatal deletion, doxycycline-containing water was administered from 8 to 10 weeks of age. *Piezo1* CKO mice were generated by crossing *Piezo1*^fl/fl^ mice with Cdh5-CreERT2 mice. Tamoxifen (75 μg/day; Sigma-Aldrich) was administered intraperitoneally once daily from postnatal day 1 to 3 (P1–P3) to induce endothelial-specific recombination. For inducible adult endothelial deletion, tamoxifen (75 μg/day, i.p.) was administered once daily for three consecutive days at 8 weeks of age. Neural crest–specific *Piezo1* deletion (*Piezo1*^ΔNC^) mice were generated by crossing *Piezo1*^fl/fl^ mice with *Wnt1*-Cre mice (B6.Cg-H2az2

Tg(*Wnt1*-cre)11Rth Tg(*Wnt1*-GAL4)11Rth/J, JAX #009107). For recombination site validation, *Piezo1*^fl/fl^; *Wnt1*-Cre and *Piezo1*^fl/fl^; Cdh5-CreERT2 mice were crossed with Rosa26mTmG reporter mice (Gt(ROSA)26Sortm4(ACTB-tdTomato,-EGFP)Luo/J, JAX #007576) to visualize Cre activity in TM and SC endothelium, respectively. *Angpt2*^fl/fl^ mice (*Angpt2*tm1c(KOMP)Seq; exon 4 floxed; MGI:5613889) were generated as previously described^63^.

*Angpt2* conditional knockout (*Angpt2* CKO) mice were generated by crossing *Angpt2*^fl/fl^ mice with Rosa26-rtTA and tetO-Cre transgenic mice; doxycycline-containing water was administered from postnatal day 0 (P0) for 2 weeks to induce recombination.

LSL-Salsa6f mice were crossed with Cdh5-CreERT2 to generate Cdh5-CreERT2^+^; LSL-Salsa6f^+/−^ reporters. To induce reporter expression, tamoxifen (75 μg/day, i.p.) was administered for three consecutive days at 8 weeks of age. Salsa6f localizes to the cytoplasm and is excluded from nuclei, enabling ratiometric intracellular Ca^2+^ measurements based on the green/red (G/R) fluorescence ratio.

*Itga9*^fl/fl^ and *Piezo1*^fl/fl^ mice were crossed with EIIa-Cre (Tg(EIIa-cre)C5379Lmgd/J, JAX #003724) mice to generate germline-deleted alleles (*Itga9*^+/-^ and *Piezo1*^+/-^), followed by Cre transgene removal by backcrossing to wild-type C57BL/6J mice. Double-heterozygous mice were obtained by intercrossing *Piezo1*^+/-^ and *Itga9*^+/-^ carriers. All strains were maintained on a mixed genetic background free from RD1 and RD8 mutations. Both male and female animals were used for all experiments. Imaging was performed at 8–10 weeks of age unless otherwise indicated. Genotyping was performed by PCR using primers listed in **Supplementary Table S1**.

### Single-cell RNA-seq re-analysis

Public mouse single-cell RNA-seq data24 were re-analyzed to compare SC endothelium and TM with respect to *Piezo1* and *Itga9* expression. Processed count matrices and the authors’ cell annotations were downloaded from the NCBI Gene Expression Omnibus (GEO Series GSE168200; BioProject PRJNA706441; SRA SRP309170). Analyses were performed in R (v4.4.2) using Seurat (v5.2.0). Cell identities followed the original annotations; all TM-related clusters reported by Thomson et al. were collapsed into a single “TM” group, and SC endothelial cluster(s) were extracted as “SC”. From this TM+SC subset, expression distributions of *Piezo1* and *Itga9* were plotted with Seurat

VlnPlot. Parameters not specified here matched the defaults in Seurat and the original study.

### Bulk RNA-seq and KEGG pathway analysis

Total RNA was isolated from LECs transfected with siControl or si*ITGA9* (harvested 48 h post-transfection; n = 6 per group) using TRIzol followed by purification with the RNeasy MinElute Kit (QIAGEN). One si*ITGA9* sample was excluded from the analysis due to insufficient RNA yield.

Library preparation and Illumina sequencing were performed by Plasmidsaurus (vendor protocol). Quality of the FASTQ files was assessed using FastQC v0.12.1. Reads were quality filtered using fastp v0.24.0 with poly-X tail trimming, 3′ quality-based tail trimming, a minimum Phred quality score of 15, and a minimum length requirement of 50 bp. Quality-filtered reads were aligned to the reference genome using STAR aligner v2.7.11 with non-canonical splice junction removal and output of unmapped reads, followed by coordinate sorting using samtools v1.22.1. PCR and optical duplicates were removed using UMI-based deduplication with UMIcollapse v1.1.0. Alignment quality metrics, strand specificity, and read distribution across genomic features were assessed using RSeQC v5.0.4 and Qualimap v2.3, with results aggregated into a comprehensive quality control report using MultiQC v1.32. Gene-level expression quantification was performed using featureCounts (subread package v2.1.1) with strand-specific counting, multi-mapping read fractional assignment, exons and 3′ UTRs as the feature identifiers, and grouping by gene_id. Final gene counts were annotated with gene biotype and other metadata extracted from the reference GTF file. Sample-sample correlations for the heatmap and PCA were calculated on normalized counts (TMM, trimmed mean of M-values) using Pearson correlation. Differential expression was performed with edgeR v4.0.16 using standard practice including filtering for low-expressed genes with edgeR::filterByExpr with default values. Differentially expressed genes (DEGs) were defined using a threshold of |log2FC| > 0.5 and FDR < 0.05. For downstream analysis, KEGG pathway enrichment analysis and bar plots were generated using the enrichR R package (v3.4).

### Outflow facility measurement

Conventional outflow facility was measured ex vivo in enucleated mouse eyes using a constant-pressure perfusion system (iPerfusion 3)^64^. Eyes were cannulated via the anterior chamber with a glass needle under a stereomicroscope and submerged in temperature-controlled PBS supplemented with 5.5 mM glucose. Eyes were then held at a physiological pressure (∼8 mmHg) for an acclimation phase of 30–45 min. After acclimation, eyes were perfused over a multi-step pressure protocol spanning the physiological range, while flow rate (Q) and pressure (P) were recorded continuously. Steady-state flow at each pressure step was identified programmatically and averaged to obtain one Q–P data point per step. The Q–P relationship was fit with a power-law model, Q(P) = Cr · P · (P/Pr)β, with Pr fixed at 8 mmHg to yield the reference outflow facility Cr (nL/min/mmHg). For ANGPT2 blockade experiments, a stopcock with a side-port was placed immediately upstream of the glass needle, and the dead volume between the side-port and the needle tip was prefilled with antibody-containing solution prior to initiating the measurement. The antibody-containing solution consisted of MEDI3617 (anti-ANGPT2, human IgG1κ; 4 mg/mL) or a matched human IgG1κ isotype control (4 mg/mL) diluted in PBS supplemented with 0.1% BSA and 5.5 mM glucose.

The acclimation phase was extended to 45–60 min to allow equilibration and delivery of antibody to the outflow pathway^65^. A paired-eye design was used such that, for each animal, one eye received MEDI3617 and the contralateral eye received isotype control, with left/right assignment randomized prior to cannulation.

### IOP measurements

IOP measurements were performed in awake mice between 9 and 11 AM using an iCare TonoLab rebound tonometer as previously described^63^. Cohorts of mutant mice and their littermate controls were measured at the indicated time points. IOP values from the left and right eyes were averaged for each animal, and these averaged values were used for all analyses reported in the manuscript.

### Endothelial Ca^2+^ imaging

Cdh5-CreERT2 mice were crossed with LSL-Salsa6f reporters, and tamoxifen (75 µg/day, i.p.) was administered for three consecutive days at 8 weeks of age to induce endothelial Salsa6f expression. Eyes were dissected to isolate a limbal strip containing SC, which was straightened on a glass slide and gently immobilized using coverslip fragments placed on both sides. Two drops of vehicle (PBS + 0.5% DMSO) or Yoda1 (20 µM) were applied to fully immerse the tissue, and time-lapse images were acquired on a Nikon Ti2 microscope for 10 min with sequential acquisition of the green (GCaMP6f) and red (tdTomato) channels using identical settings across conditions. Images were analyzed in Fiji/ImageJ by defining an SC endothelial ROI on the tdTomato channel and applying it to both channels across all frames; background-subtracted mean intensities were measured per frame and Ca^2+^ responses were quantified as the ratiometric green/tdTomato signal, G/R = (G − Gbg)/(R − Rbg), with optional normalization to baseline (t0).

### Validation of *Itga9* exon 9 excision by Sanger sequencing

To confirm Cre-mediated excision of *Itga9* exon 9, total RNA was isolated from lung using TRIzol (15596018; Life Technologies) following the standard extraction protocol. cDNA was generated using the iSCRIPT synthesis kit (1708891; Bio-Rad) according to the manufacturer’s instructions. PCR amplification across the exon 9 region was performed using cDNA as template (**Supplementary Table S1**). PCR products were purified using the Wizard SV Gel and PCR Clean-Up System (Promega, A9281) and submitted to ACGT DNA Services for Sanger sequencing.

### Cell culture and Yoda1 stimulation

Primary HDLECs (PromoCell, C-12216) were cultured in endothelial cell growth medium MV2 (PromoCell, C-22022) supplemented with the manufacturer’s supplement mix (SupplementMix; PromoCell, C-39226), in glass-bottomed culture plates at 37 °C in a humidified incubator with 5% CO2. Cells between passages 3 and 6 were used for all experiments. For experiments requiring serum starvation, cells were incubated overnight in Endothelial Cell Basal Medium (PromoCell, C-22210) prior to stimulation. For siRNA-mediated knockdown, HDLECs were transfected with siRNAs targeting *ANGPT2* (Ang-2 siRNA (h); Santa Cruz Biotechnology, sc-39305; final concentration 15 nM) or a non-targeting control siRNA (Control siRNA-A; Santa Cruz Biotechnology, sc-37007) using Lipofectamine RNAiMAX Transfection Reagent (Thermo Fisher Scientific/Invitrogen; Cat. No. 13778030) according to the manufacturer’s instructions. Cells were incubated for 48 hours after transfection before downstream experiments. For pharmacological stimulation, HDLECs were treated with 0.5 μM Yoda1 (MilliporeSigma, Cat. No. SML1558-5MG) or vehicle control (DMSO) for 5 minutes at 37 °C prior to fixation.

### Immunofluorescence staining

Cells or tissues were fixed in 4% paraformaldehyde (PFA) for 10 minutes at room temperature, permeabilized with TBS containing 0.1% Triton X-100 for 10 minutes and blocked with TBS containing 5% donkey serum for 1 hour at room temperature. Primary and secondary antibodies were diluted in the same blocking buffer and incubated at room temperature. Nuclei were counterstained with DAPI. Fluorescence images were acquired using a Nikon A1 confocal microscope. All image processing and analysis were performed using Fiji software (ImageJ 1.54p; Java 1.8.0_322, 64-bit). Antibody information is provided in **Supplementary Table S2**. For colocalization analysis, Pearson’s correlation coefficient (R) was calculated between the red (ZO-1) and green (ITGA9 or ITGB1) channels using the Coloc2 plugin in Fiji with the “above threshold” setting to reduce background signal. At least 12 images (40× objective) per group were acquired across three independent experiments for analysis.

### VE-cadherin junction width quantification

VE-cadherin junction widths were quantified from VE-cadherin immunofluorescence images using a previously described custom Python script^66^. Briefly, VE-cadherin–positive junctions were segmented from background by applying an intensity threshold to generate a binary mask. The program randomly sampled points within the VE-cadherin–positive regions and calculated junction width by determining the shortest path across the junction structure through each point, defined as the minimum sum of distances to the first VE-cadherin–negative regions on either side. Ten fields of view were randomly chosen and masked from each sample for quantification.

### Western blotting

HDLECs were transfected with siRNAs using Lipofectamine RNAiMAX reagent (Thermo Fisher Scientific, 13778075) according to the manufacturer’s instructions. Integrin α9/*ITGA9* siRNA (h) (Santa Cruz Biotechnology, sc-75340) and Ang-2 siRNA (h) (Santa Cruz Biotechnology, sc-39305) and Silencer Select siRNA targeting human *TEK* (TIE2) (Assay ID s13983; Thermo Fisher Scientific, Ambion; Cat# 4390825) were used, along with the respective negative control siRNAs (Control siRNA-A; Santa Cruz Biotechnology, sc-37007 and Silencer™ Negative Control No. 1 siRNA; Thermo Fisher Scientific; Cat# AM4635), at a final concentration of 15 nM. Transfection was performed on subconfluent cultures, and cells were harvested for downstream analysis 48 hours after transfection. For experiments requiring serum starvation, cells were incubated overnight in Endothelial Cell Basal Medium (PromoCell, C-22210) prior to stimulation. For Yoda1 stimulation, 0.5 μM Yoda1 or DMSO was added 5 minutes at 37 °C prior to cell lysis. For recombinant ANGPT2 stimulation, cells were treated with recombinant human ANGPT2 (Acro Biosystems, Cat. No. AN2-H5242) at 300 or 900 ng/mL (vehicle: PBS) for 30 minutes prior to cell lysis. For ANGPT2 neutralization, cells were incubated with MEDI3617 (Novus Biologicals, Cat. No. NB465096) or matched human IgG1κ isotype control (Cat. No. 0151K-14) at 2 μg/mL for 4 hours prior to Yoda1 stimulation; antibodies were present during Yoda1 stimulation.

HDLECs and tissues from *Itga9* WT or CKO mice were lysed with RIPA buffer supplemented with Halt protease and phosphatase inhibitor cocktails (78442; Thermo Fisher Scientific) on ice for 30 min, and protein concentration was measured by a BCA assay (Thermo Fisher Scientific). Equal amounts of protein were mixed with Laemmli sample buffer (1% SDS, 62.5 mM Tris-HCl [pH 6.8], 10% glycerol, 5% β-mercaptoethanol, 0.05% bromophenol blue), boiled for 10 min, separated by SDS-PAGE, and transferred to PVDF membranes (GE Healthcare). Membranes were blocked with 5% BSA in TBST (Tris-buffered saline with 0.1% Tween-20) for 1 h at room temperature, incubated with primary antibodies overnight at 4°C, washed three times with TBST, and incubated with appropriate HRP-conjugated secondary antibodies for 1 h at room temperature. Antibody information used for immunoblotting is provided in **Supplementary Table S2**. Chemiluminescent signal was developed using an ECL substrate and imaged using an iBright Imaging System (iBright 1500; Thermo Fisher Scientific).

### OCT acquisition and peripapillary retinal nerve fiber layer thickness (pRNFLT) measurement

In vivo OCT was performed on a Heidelberg Spectralis (HRA+OCT/OCT2) equipped with the animal imaging stage and the mouse adapter lens +25 D (Heidelberg Engineering). Mice were anesthetized by intraperitoneal ketamine/xylazine for OCT imaging. The optic nerve head (ONH) was centered on the infrared fundus image, and scans were acquired in a radial pattern.

Exported B-scans (TIFF) were analyzed in ImageJ. Images were calibrated using the scale bar embedded in each exported image (µm/pixel). Retinal nerve fiber layer thickness (RNFLT) was measured on each B-scan as the perpendicular distance from the inner limiting membrane to the outer RNFL boundary. Measurement sites were placed 200 µm from the ONH center along the nasal, temporal, superior, and inferior meridians (one point each)^67^. For each eye, the four values were averaged to yield an eye-level peripapillary RNFLT (pRNFLT); for each mouse, the average of both eyes (eight measurements total) provided one pRNFLT value.

### Intracameral injection and fixation

Mice were anesthetized with isoflurane (3% for induction, 1.5–2% for maintenance) and placed on a heated platform to maintain body temperature. FluoSpheres™ (20 nm, carboxylate-modified polystyrene, Thermo Fisher Scientific) were diluted in sterile PBS containing Ca^2+^/Mg^2+^ to a final concentration of 1 × 10^11^ particles/mL (∼0.06% v/v). A total of 1 μL of tracer solution was sequentially loaded into a 10 μL NanoFil syringe (World Precision Instruments) and separated by a small air bubble (∼0.2 μL). The syringe was fitted with a 35G beveled needle (NF35BV, WPI) and mounted on an UltraMicroPump (UMP3, WPI) connected to a MICRO-2T SMARTouch™ controller (WPI) and a micromanipulator. The tracer was first delivered into the anterior chamber. Yoda1 (20 μM in PBS containing 0.5% DMSO) was injected in the same manner as described above. Eyes were collected 30 min after injection and immersion-fixed overnight in 2% paraformaldehyde (PFA) at 4 °C.

### Transmission electron microscopy (TEM)

For TEM analysis, mice were anesthetized and transcardially perfusion-fixed with 4% paraformaldehyde and 2% glutaraldehyde in 0.1 M sodium cacodylate buffer (pH 7.2). Eyes were then enucleated and post-fixed overnight by immersion in the same fixative at 4 °C. Samples were transferred as whole globes to the Gong laboratory (Boston University) for TEM processing and imaging (Hongyuan Ren and Haiyan Gong). Anterior segments were carefully dissected into radial wedges and sequentially dehydrated in graded ethanol solutions (50%, 70%, 80%, 95%, and 100%). After washing in propylene oxide, samples were infiltrated with a 1:1 mixture of propylene oxide and Epon–Araldite resin for 3 h, followed by infiltration with 100% Epon–Araldite overnight and embedding in fresh resin. Semi-thin sections (4 µm) were cut and stained with 1% toluidine blue to identify the aqueous humor outflow pathway by light microscopy. Ultrathin sections (70 nm) were then cut and imaged using a transmission electron microscope (JEOL JEM-1400 Flash) at 1,000× magnification. Serial TEM images were acquired along the inner wall of SC and digitally stitched to generate mosaic views for each section. Using ImageJ, SC inner-wall length was measured against the scale bar in each stitched image, and GVs were counted. GV number was normalized to 100 μm of SC inner wall for each section.

### EdU assay

To label proliferating cells, EdU (5-ethynyl-2’-deoxyuridine; Carbosynth) was administered in the drinking water at a final concentration of 200 μg/mL for 7 consecutive days. EdU was prepared as a 10 mg/mL stock solution in DMSO and diluted 1:50 in sterile drinking water. The solution was replaced every 2 days and protected from light using foil-wrapped bottles. After the labeling period, mice were euthanized and the anterior segment of the eyes were carefully dissected and fixed in 2% PFA in PBS overnight at 4 °C. The next day, tissues were washed in TBS-T (3 × 5 min) and permeabilized/blocking buffer was applied for 1 hour at room temperature. The blocking buffer consisted of 5% donkey serum, 2.5% bovine serum albumin (BSA), and 0.5% Triton X-100 in TBS. EdU detection was performed using a custom copper-catalyzed azide–alkyne cycloaddition (CuAAC) reaction. A Click reaction buffer was freshly prepared containing 4 mM CuSO4 (Acros Organics), 100 mM sodium ascorbate (Acros Organics; freshly made), and 5 μM sulfonated Alexa Fluor azide (e.g., Sulfo-Cyanine3 Azide, Lumiprobe) in TBS (pH 7.6). Tissues were incubated in this buffer for 30–60 minutes at room temperature in the dark, followed by 3 washes in PBS (10–15 min each). Sulfonated azide dyes were used to reduce non-specific background staining in whole-mount tissue. Following EdU development, tissues were again blocked for 15–30 minutes and subjected to immunofluorescence staining. Samples were incubated with primary antibodies diluted in TBS containing 1% BSA and 0.3% Triton X-100, overnight at 4 °C. After washing (3 × 15 min in TBS), tissues were incubated with species-appropriate fluorescent secondary antibodies for 1 hour at room temperature in the dark. Nuclei were counterstained with DAPI (1 μg/mL, 15 min to overnight), followed by a final wash and mounting with antifade medium. All steps involving fluorophores were carried out under light-protected conditions.

### SC and RGC immunofluorescence imaging and quantification

To evaluate SC and RGCs, whole-mounted anterior segments and retinas were subjected to immunofluorescence staining. Enucleated eyes were fixed overnight in 2% paraformaldehyde (PFA) at 4 °C. After fixation, conjunctiva and residual connective tissue were removed. A circumferential incision was made approximately 1 mm posterior to the limbus to remove the posterior segment and lens, followed by four radial incisions to prepare a petal-shaped anterior flat mount. Tissues were blocked and permeabilized for 1 hour at room temperature in TBS containing 5% donkey serum, 1% BSA, and 0.3% Triton X-100. Samples were incubated overnight at 4 °C with primary antibodies, followed by TBS-T washes and detection with species-appropriate fluorophore-conjugated secondary antibodies. Primary and secondary antibodies used are listed in **Supplementary Table S2**. After immunostaining, small relaxing cuts were made around the cornea to flatten the SC region. The tissue was mounted in antifade reagent with the outer scleral surface facing the coverslip. Confocal imaging was performed using a Nikon A1 microscope to acquire Z stacks of the central region of each quadrant using a 20× objective lens. Maximum-intensity projections were used for quantification, and canal area was measured in Fiji (ImageJ 1.54p; Java 1.8.0_322, 64-bit). Expression levels within SC (e.g., p-AKT, ITGA9, p-FAK) were reported as background-subtracted mean fluorescence per mm^2^ of CD31^+^ SC area. Background subtraction was performed by subtracting the mean gray value of a same-field cell-free region of interest (ROI) from the mean gray value within the SC ROI; negative values were clipped to zero. All acquisition settings (laser power, detector gain, offset, pinhole, pixel size, and Z-step) were kept identical across groups. All image analysis was performed in a blinded fashion. For RGC analysis, retinas were flat-mounted and imaged using a Ti2 microscope (Nikon) at 20× magnification. Four images each were obtained from the central (0.1 mm from the optic disc), middle (0.8 mm), and peripheral (1.5 mm) retina in each quadrant (total of 12 fields per retina). Images were cropped to 200 × 200 μm and RGCs were manually counted. The average RGC density (cells/mm^2^) was calculated.

### Statistical analysis

Analysis of physiological, histological, and single-cell transcriptomic data was performed using GraphPad Prism 10.0.4 (GraphPad Software, San Diego, CA, USA), R version 4.4.2, or JMP version 16.0.0 (SAS Institute, Cary, NC, USA). Statistical significance was assessed using unpaired two-tailed Student’s t-test or one-way ANOVA with Tukey–Kramer’s test, as appropriate. The specific statistical test used for each dataset is indicated in the corresponding figure legend. All data are presented as mean ± SEM. Outflow facility Cr is approximately log-normally distributed; therefore, group comparisons were performed using natural log-transformed values, ln(C_r_)^64^. For two-group comparisons (*Itga9* WT vs *Itga9* CKO; *Piezo1* WT vs *Piezo1*^ΔEC^), ln(C_r_) was analyzed using an unpaired two-tailed Student’s t-test. A two-sided P < 0.05 was considered statistically significant. P values less than 0.05 were considered statistically significant and are denoted as follows: *P < 0.05, **P < 0.01, ***P < 0.001, ****P < 0.0001. All analyses were performed in a blinded fashion with respect to treatment or genotype.

## Data Availability

The data that support the findings of this study are available from the corresponding author upon reasonable request. Public single-cell RNA-seq data used for re-analysis are available from the NCBI Gene Expression Omnibus (GEO) under accession GSE168200.

## Acknowledgements

Confocal imaging was performed at the Center for Advanced Microscopy of the Feinberg School of Medicine, supported by NCI CCSG P30 CA060553.

Electron microscopy was performed at the Transmission Electron Microscope Core at Boston University Chobanian & Avedisian School of Medicine, supported by NIH S10OD028571. We thank Dr. Livingston Van De Water (Albany Medical College) for providing the Itga9fl/fl mice and Dr. Gou Young Koh (KAIST) for generously sharing the anti-ANGPT2 antibody. We also thank Dr. Gregory W. Schwartz (Feinberg School of Medicine, Northwestern University) for providing access to the LSL-Salsa6f transgenic line (ratiometric tdTomato–GCaMP6f; JAX #031968). We thank Joseph van Batenburg-Sherwood for setting up the iPerfusion outflow facility system used in this study.

Mouse illustrations in Figs. 1B, 1I, 3C, 5B and 5K were created using BioRender.com.

