## Supplementary information for "Piezo1 Triggers an Angiopoietin-2-Integrin Signaling Loop in Schlemm’s Canal to Regulate Intraocular Pressure"

676 N. St. Clair Ave., Suite 2300, Chicago, IL 60611, USA

### Supplementary Figures

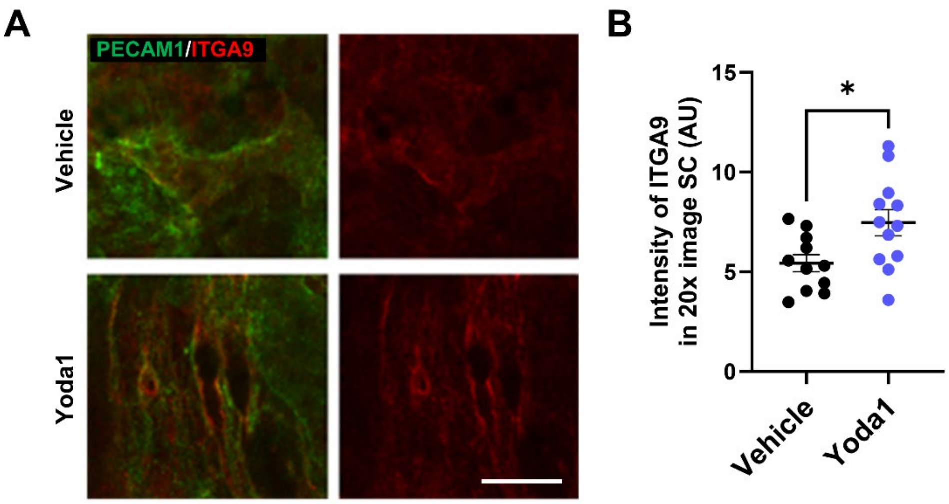


**Supplementary Figure S1. Yoda1 increases ITGA9 immunoreactivity in SC.**

(A) Representative images of SC stained for PECAM1 (green) and ITGA9 (red) in vehicle- and Yoda1-treated samples; the ITGA9 single channel is shown at right. Scale bar, 100 μm. (B) Quantification of ITGA9 fluorescence intensity in SC per 20× image (AU). n = 11–12 eyes per group. Bars indicate mean ± SEM. Statistics: unpaired two-tailed Student's t-test; $P = 0.0179$. $P < 0.05$.

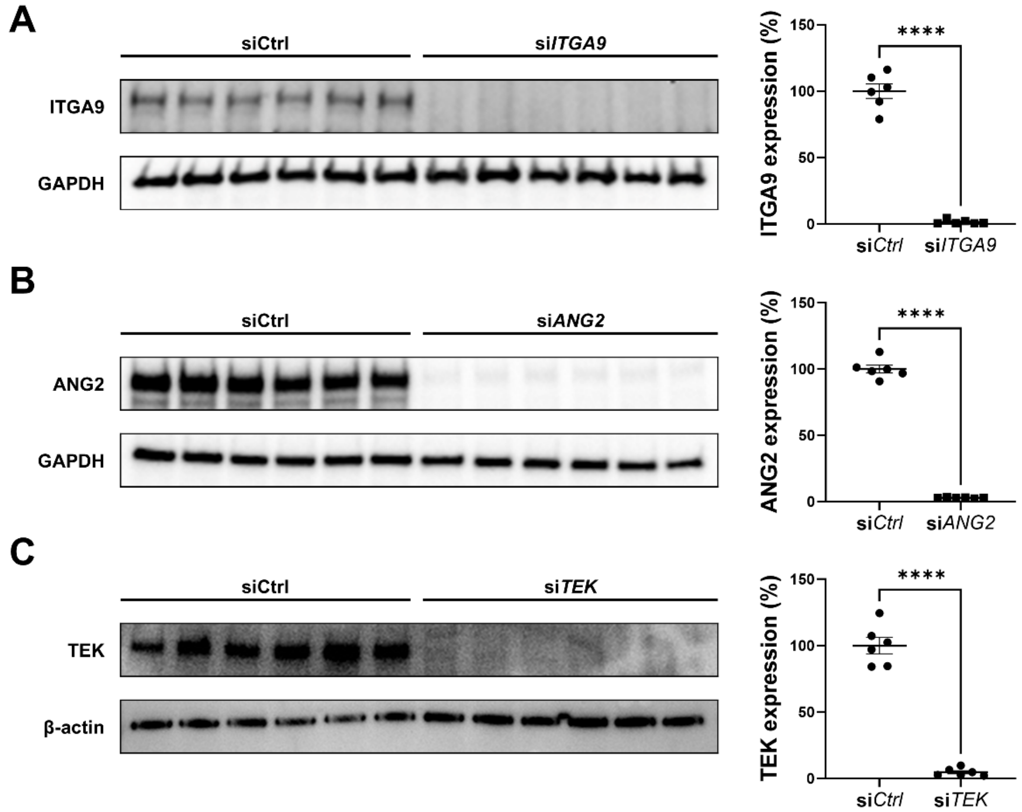


**Supplementary Figure S2. Efficient siRNA-mediated knockdown of *ITGA9* and *ANGPT2* (ANG2) in HDLECs.**

(A) Representative immunoblot of *ITGA9* and GAPDH from HDLECs transfected with control siRNA (siCtrl) or si*ITGA9* (n = 6 per group). Densitometric quantification of *ITGA9* normalized to GAPDH and expressed as percent of siCtrl confirmed robust *ITGA9* depletion (98.4% reduction). (B) Representative immunoblot of *ANGPT2* (ANG2) and GAPDH from HDLECs transfected with siCtrl or siANG2 (n = 6 per group). Densitometric quantification of ANG2 normalized to GAPDH and expressed as percent of siCtrl confirmed robust ANG2 depletion (97.0% reduction). (C) Representative immunoblot of *TEK* and β-actin from HDLECs transfected with siCtrl or si*TEK* (n = 6 per group). Densitometric quantification of *TEK* normalized to β-actin and expressed as percent of siCtrl confirmed robust *TEK* depletion (95.2% reduction).

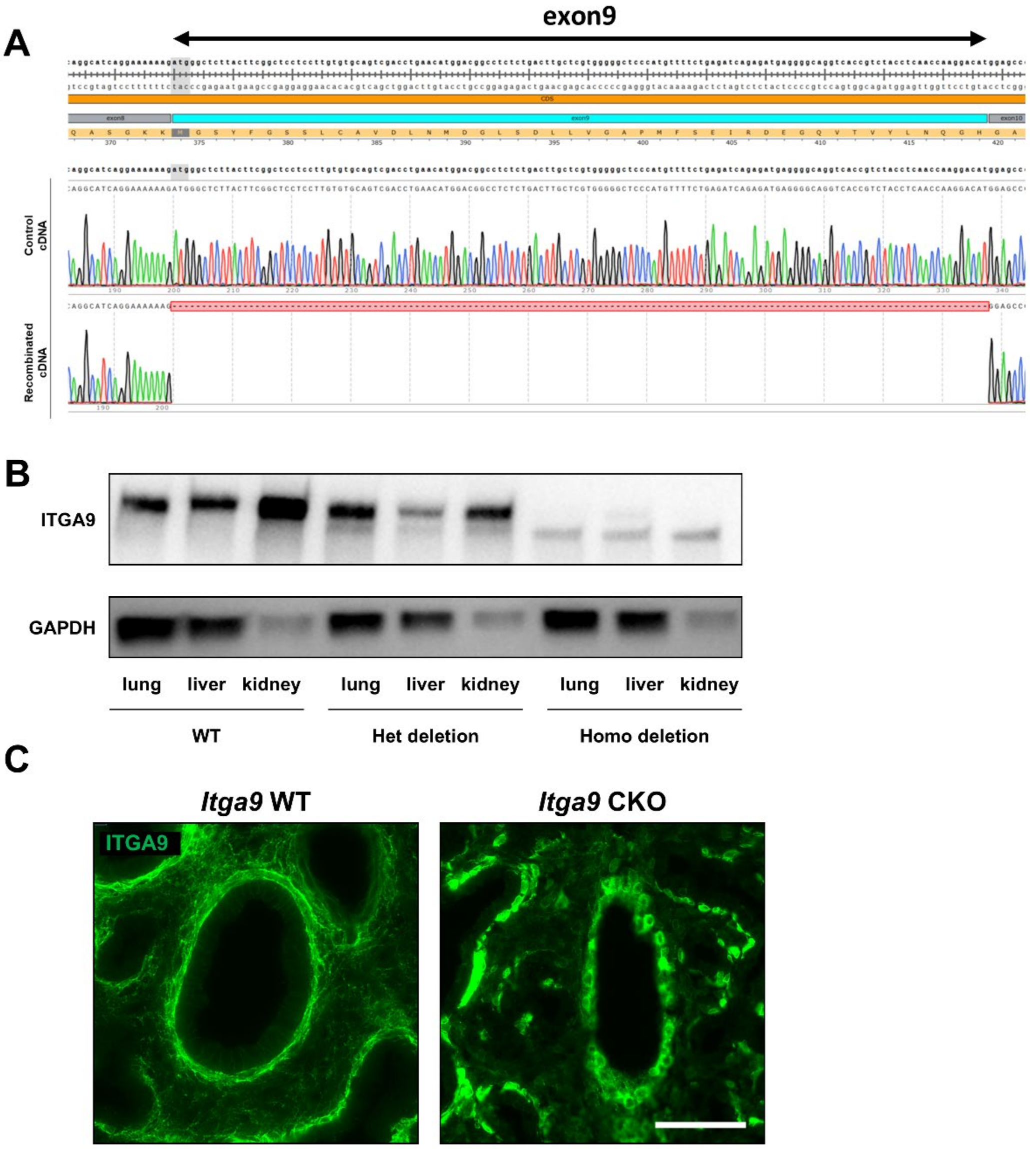


**Supplementary Figure S3. Cre-mediated excision of *Itga9* exon 9 produces a residual truncated ITGA9 species and alters ITGA9 subcellular localization in *Itga9* CKO mice.**

(A) Sanger sequencing of RT-PCR amplicons spanning the *Itga9* exon 9 region shows exon 9 skipping (exon 8–exon 10 junction) in recombined cDNA compared with control. (B) Immunoblotting of ITGA9 in lung, liver, and kidney lysates from *Itga9* WT, heterozygous deletion, and homozygous deletion mice; GAPDH serves as a loading control. Full-length ITGA9 is markedly reduced in homozygous deletion tissues, with an additional lower-molecular-weight immunoreactive band consistent with a residual truncated ITGA9 species. (C) Representative ITGA9 immunofluorescence in lung sections from *Itga9* WT (left) and *Itga9* CKO (right) mice, showing redistribution from a membrane-associated pattern to a predominantly intracellular pattern in recombined samples. Images were acquired and displayed using identical settings. Scale bar, 50 μm. n = 3 mice per group.

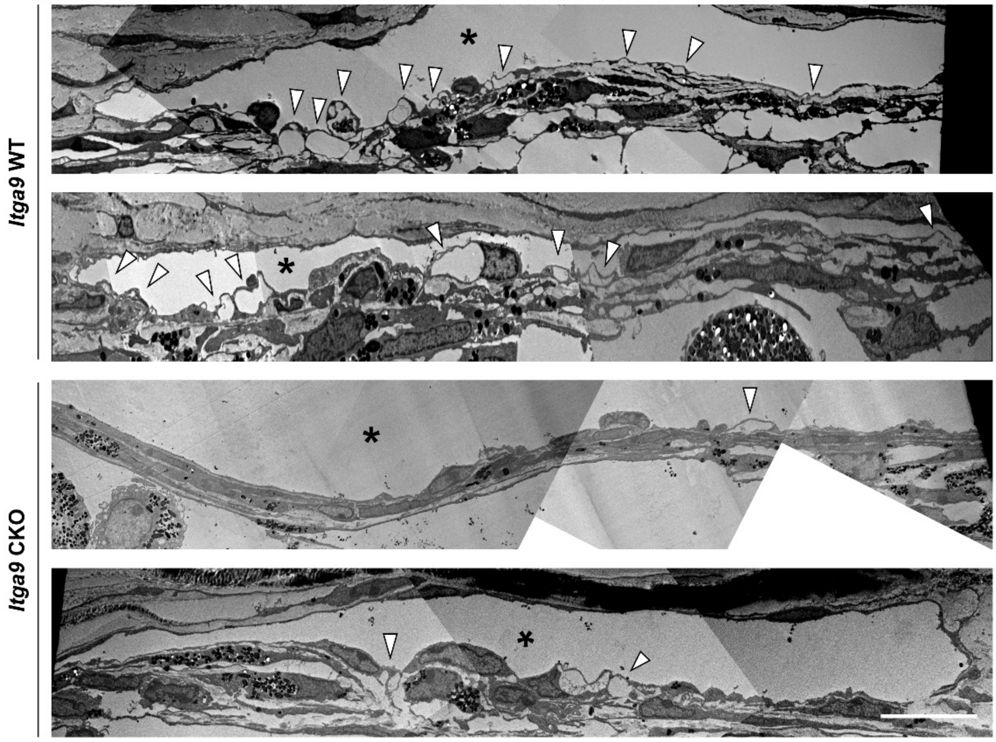


**Supplementary Figure S4. Ultrastructural appearance of the Schlemm's canal inner wall in *Itga9* WT and *Itga9* CKO eyes.**

Representative stitched transmission electron microscopy (TEM) mosaics showing the Schlemm's canal (SC) inner wall in *Itga9* WT and *Itga9* CKO eyes. Two representative sections from each genotype are shown. Arrowheads indicate giant vacuoles (GVs) along the SC inner wall, and asterisks indicate the SC lumen. GVs were less prominent in *Itga9* CKO eyes than in WT eyes. Images are shown with the scleral side oriented upward. Scale bar, 10 μm.

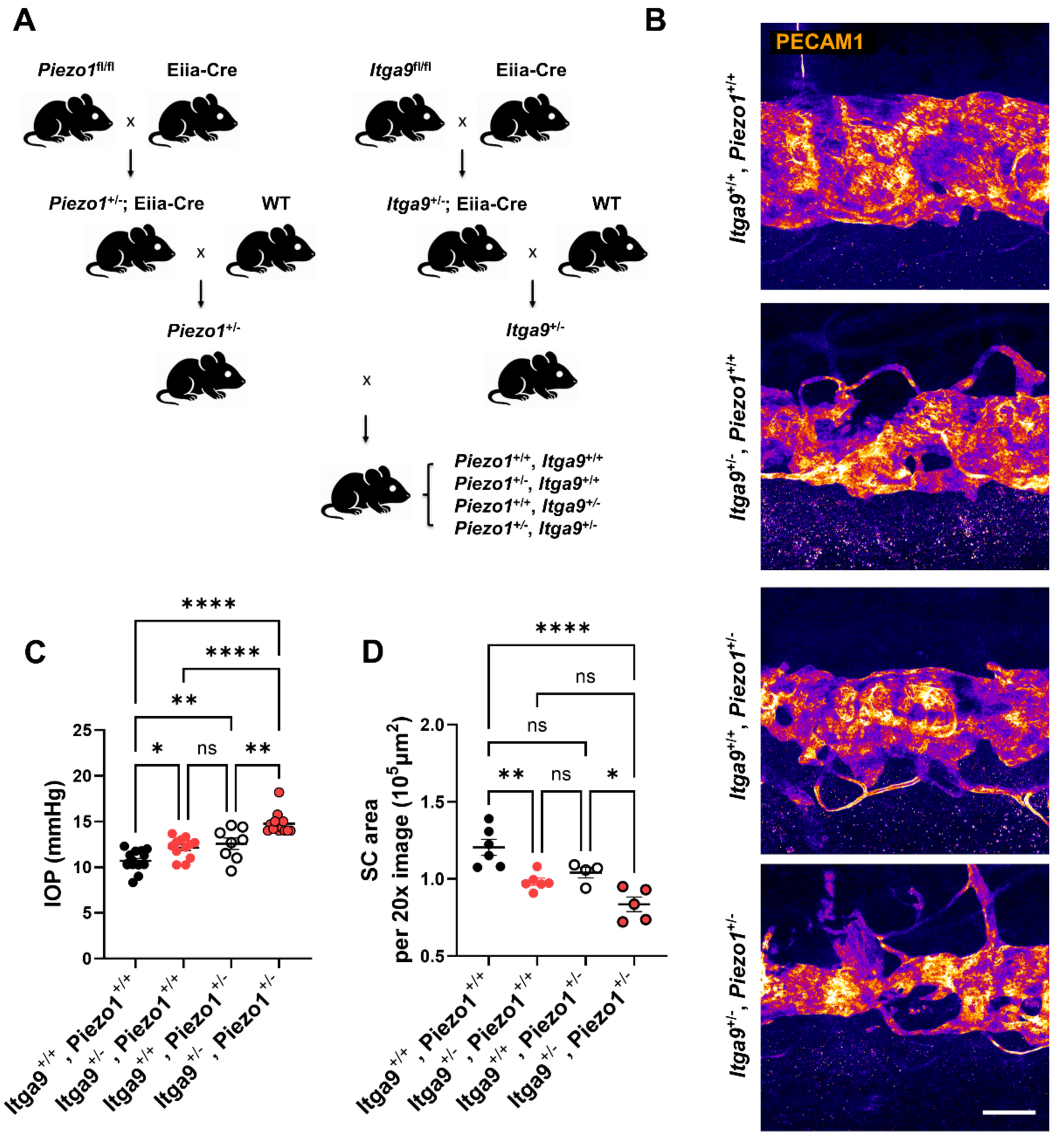


**Supplementary Figure S5. Combined heterozygous deletion of *Piezo1* and *Itga9* impairs SC morphology and elevates IOP.**

(A) Breeding scheme for generating *Piezo1*$^{+/-}$; *Itga9*$^{+/-}$ double heterozygous mice. Germline deletion was induced by crossing *Piezo1*$^{fl/fl}$ or *Itga9*$^{fl/fl}$ mice with Eiia-Cre, and Cre was subsequently bred out by crossing with WT mice. The resulting *Piezo1*$^{+/-}$ and *Itga9*$^{+/-}$ mice were intercrossed to generate four genotypes: *Piezo1*$^{+/+}$; *Itga9*$^{+/+}$, *Piezo1*$^{+/-}$; *Itga9*$^{+/+}$, *Piezo1*$^{+/+}$; *Itga9*$^{+/-}$, and *Piezo1*$^{+/-}$; *Itga9*$^{+/-}$. (B) Representative SC whole-mount images stained with PECAM1 at 8 weeks of age for each genotype. (C) IOP measurements showed significant increases in both single heterozygotes versus WT, and were further elevated in *Piezo1*$^{+/-}$; *Itga9*$^{+/-}$ mice compared with either single heterozygote and WT. (D) SC area was significantly reduced in *Itga9*$^{+/-}$; *Piezo1*$^{+/+}$ mice, showed a trend toward reduction in *Piezo1*$^{+/-}$; *Itga9*$^{+/+}$ mice, and was further decreased in the double heterozygotes compared with WT and *Piezo1*$^{+/-}$; *Itga9*$^{+/+}$ mice (with a trend versus *Itga9*$^{+/-}$; *Piezo1*$^{+/+}$ mice). Data are presented as mean ± SEM. Statistical analysis was performed using one-way ANOVA followed by Tukey–Kramer post hoc test. *P < 0.05, **P < 0.01, ***P < 0.001, ns = not significant. Scale bar: (B) 100 μm.

**Table S1. Primer sequences used for genotyping**

| Primer set / Target | Primer | Sequence (5'→3') |
|---|---|---|
| *Itga9* floxed allele | forward | CCTTACAGGGCTCTAGGAAAGGGG |
| | reverse | AATAGTCATTGAGACTCTCCCTGG |
| *Itga9* exon 9 excision (Sanger) | forward | GCTGAACCTCACGGACAACA |
| | reverse | GTCAGGGTCAGCTGTTCCTC |
| *Piezo1* floxed allele | forward | GCCTAGATTCACCTGGCTTC |
| | reverse | GCTTTAACCATTGAGCCATCT |
| mTmG (GFP cassette) | forward | GGGCACAAGCTGGAGTACAA |
| | reverse | GTCCATGCCGAGAGTGATCC |
| Rosa26-rtTA allele | forward | AAGGGAGCTGCAGTGGAGTA |
| | Mutant reverse | GGCGAGTTTACGGGTTGTTA |
| | WT reverse | TCCGAGGCGGATCACAAGCA |
| Generic Cre | forward | GTGCAAGTTGAATAACCGGAAATGG |
| | reverse | AGAGTCATCCTTAGCGCCGTAAATCAAT |
| Cdh5-CreERT2 | forward | ATGCAAGCTGGTGGCTGGACC |
| | reverse | GATCTCCACCATGCCCTCTACAC |
| *Angpt2* floxed allele | forward | GGGAAACCTCAACACTCCAA |
| | reverse | ACACCGGCCTCTAGACACAC |
| Rosa26-LSL-Salsa6f | WT forward | CTGGCTTCTGAGGACCG |
| | Mutant reverse | TGGTAGTGGTAGGCGAGCTG |
| | WT reverse | CCGAAAATCTGTGGGAAGTC |
| | Mutant forward | CTGTTCCTGTACGGCATGG |

**Table S2. Materials used in the present study**

| Reagent | Company | Identifiers |
|---|---|---|
| Anti-PECAM-1 (CD31) | BD Biosciences | 553370 |
| Anti-ITGA9 | R&D Systems | AF3827 |
| Anti-ITGB1 | R&D Systems | MAB17781 |
| Anti-ITGB1 (active conformation; clone 9EG7) | BD Biosciences | 550531 |
| Anti-ZO-1 | Invitrogen | 40-2200 |
| Anti-Phospho-FAK (Tyr397) | Invitrogen | 44-625G |
| Anti-FAK (total) | Cell Signaling Technology | 3285S |
| Anti-Phospho-TIE2 | R&D Systems | AF2720 |
| Anti-TIE2/TEK (total; clone Ab33) | MilliporeSigma | 05-584 |
| Anti-Phospho-AKT (Ser473) | Cell Signaling Technology | 9271 |
| Anti-FOXO1 (clone: C29H4) | Cell Signaling Technology | 2880S |
| Anti-RBPMS | PhosphoSolutions | 1832-RBPMS |
| Anti-PROX1 | R&D Systems | AF2727 |
| Anti-Integrin α9 (Human) | R&D Systems | MAB4574 |
| ERG | Abcam | AB92513 |
| Anti-Human VE-Cadherin | R&D Systems | AF938 |
| GAPDH (14C10) | Cell Signaling Technology | 2118S |
| Anti-α Tubulin (DM1A) | Santa Cruz Biotechnology | sc-32293 |
| Anti-ANGPT2 | Gift from Dr. Gou Young Koh | NA |
